# barCoder: a tool to generate unique, orthogonal genetic tags for qPCR detection

**DOI:** 10.1101/2020.01.06.896035

**Authors:** Casey B. Bernhards, Matthew W. Lux, Sarah E. Katoski, Tyler D. P. Goralski, Alvin T. Liem, Henry S. Gibbons

**Author notes:** These authors contributed equally to this work (Casey B. Bernhards, Matthew W. Lux).

## Abstract

**Background:** Tracking dispersal of microbial populations in the environment requires specific detection methods that discriminate between the target strain and all potential natural and artificial interferents, including previously utilized tester strains. Recent work has shown that genomic insertion of short identification tags, called “barcodes” here, allows detection of chromosomally tagged strains by real-time PCR. Manual design of these barcodes is feasible for small sets, but expansion of the technique to larger pools of distinct and well-functioning assays would be significantly aided by software-guided design.

**Results:** Here we introduce barCoder, a bioinformatics tool that facilitates the process of creating sets of uniquely identifiable barcoded strains. barCoder utilizes the genomic sequence of the target strain and a set of user-specified PCR parameters to generate a list of suggested barcode “modules” that consist of binding sites for primers and probes and appropriate spacer sequences. Each module is designed to yield optimal PCR amplification and unique identification. Optimal amplification includes metrics such as ideal T_m_ and G+C-content, appropriate spacing, and minimal stem-loop formation; unique identification includes low BLAST hits against the target organism, previously generated barcode modules, and databases, such as NCBI. We tested the ability of our algorithm to suggest appropriate barcodes by generating 12 modules for *Bacillus thuringiensis* serovar *kurstaki*, a simulant for the potential biowarfare agent *Bacillus anthracis*, and three each for other potential target organisms with variable G+C content. Real-time PCR detection assays directed at barcodes were specific and yielded minimal cross-reactivity with a panel of near-neighbor and potential contaminant materials.

**Conclusions:** The barCoder algorithm facilitates the generation of barcoded biological simulants by (a) eliminating the task of creating modules by hand, (b) minimizing optimization of PCR assays, and (c) reducing effort wasted on non-unique barcode modules.

## Background

Developing an understanding of organisms in their natural ecological niches requires the ability to measure the dynamic interaction with their environment, either at the level of the individual or at population scales. For metazoa, a number of approaches have been utilized to track individuals of a species, including simple bands or markings conferring unique identifiers, Passive Integrated Transponders (PITs), telemetry devices, and biologgers (1, 2). These approaches are limited to large organisms, as they require either direct visual inspection or electronic devices that can be attached by physical means to the body of the organism in question. As the field of environmental microbiology continues to mature, novel tools to facilitate “tag and release” studies are critical to understanding microbial interactions within existing environmental niches or in the context of introduction into new environments. Early efforts to track environmental fate of genetically modified organisms in field releases utilized fluorescently or metabolically marked strains of *Pseudomonas putida* (3, 4) and *P. fluorescens* (5, 6). Likewise, spontaneous rifampicin-resistant mutants have been used to track establishment and persistence of introduced isolates in field trials (7). However, conventional selectable, chromogenic, or fluorescent markers carry metabolic costs that can compromise the carrier strain’s fitness in resource-constrained environments (8), revealing the need for phenotypically neutral, non-coding, genomic insertions that can differentiate introduced strains from native flora.

The development of DNA synthesis chemistry, microarray technology, quantitative PCR (qPCR) and high-throughput sequencing resulted in the development of several important capabilities and tagging approaches. Early studies used transposons containing short synthetic barcodes to identify virulence factors in several organisms (9, 10). As oligonucleotide synthesis technology became more sophisticated and costs decreased, longer tags could be produced, resulting in the use of tagged strains to study the spatiotemporal dispersion in systems otherwise unamenable to tracking. In particular, significant work has been done to understand the details of stochastic dynamics of *Salmonella* infections by monitoring the relative quantities of tagged strains in different locations within the host (11-13). These tagged strains, known as Wild-type Isogenic Tagged Strains (WITS), contain short, unique sequences inserted into the genome to allow quantitation by qPCR (11). Similar work has been done to study population dynamics during infection for several other bacterial and viral pathogens (13-20).

The ability to track the fate of microbes introduced into an environment is also of interest to the biodefense research community. Spores of *Bacillus anthracis*, the causative agent of anthrax, were used in the high-profile 2001 anthrax mail attacks and were historically weaponized by both the United States and Russia on large scales (21). An important angle for preparedness against a potential attack includes an understanding of how spores released into the environment might disperse, persist, and migrate. The release of live *B. anthracis* spores (and indeed, even of attenuated strains) in an outdoor test is impossible due to public health concerns. Instead, close biological relatives are used as simulants. In the case of *B. anthracis*, recent work has used *Bacillus thuringiensis* serovar *kurstaki* (Btk) due to its similar physiological and biochemical properties (22-26). Yet, even with the use of an adequate simulant, repeated dispersion testing on the same test site is problematic due to a need to distinguish between past and present testing, especially for a ubiquitous environmental bacterium such as Btk that is also in widespread use as a commercial biopesticide (27-30). In addition, the problem of “signature erosion” has diminished the utility of endogenous genomic signatures as detection tools as the diversity of sequence data in public databases has exploded (31, 32).

To overcome these challenges, we previously inserted unique genetic barcodes designed to enable rapid detection by qPCR into the Btk genome (24) and subsequently tested the system in a field release (23). These strains were successfully detected in field samples using the qPCR assays, but, like the earlier WITS strains, the strains constructed for our field release (23) did not exploit the full ability of bioinformatics and synthetic biology that has become available. Most notably, each of the tags required its own specific PCR assay conditions, which makes scaling up to larger numbers of barcodes prohibitive. In this work, we have built upon our previous work by developing an algorithm, called barCoder, to generate barcode sequences that are unique amongst a pool of barcoded strains and require minimal development of qPCR assays. The algorithm also provides numerous features to minimize experimental troubleshooting efforts and customize amplicon properties. Here, we present the algorithm, as well as experimental validation of its ability to generate a potentially unlimited pool of highly diverse DNA barcodes, each with its own specific qPCR assay.

## Results

### Barcode design

Two major types of qPCR assays exist: assays based on intercalating dyes (e.g. SYBR Green) responsive only to double-stranded DNA, and assays based on 3□-exonuclease degradation of probe sequences effecting a signal unquenching (referred to herein as TaqMan for the probes used). In terms of assay design, both require a forward and reverse primer with similar constraints such as amplicon length, T_m_, G+C-content, G+C-clamp, potential secondary structure, and primer dimer formation. The primary design difference between the qPCR assay types is a third probe sequence required only for TaqMan assays with its own recommended design guidelines. Thus, in general, the same primer set can be used for either approach with the same barcode (Fig 1A). From the perspective of qPCR assay design, the remaining spacer sequences between primers/probe are largely immaterial other than to meet ideal amplicon size targets, and therefore can be generated randomly, with constraints (see Algorithm design).

**Figure 1.**
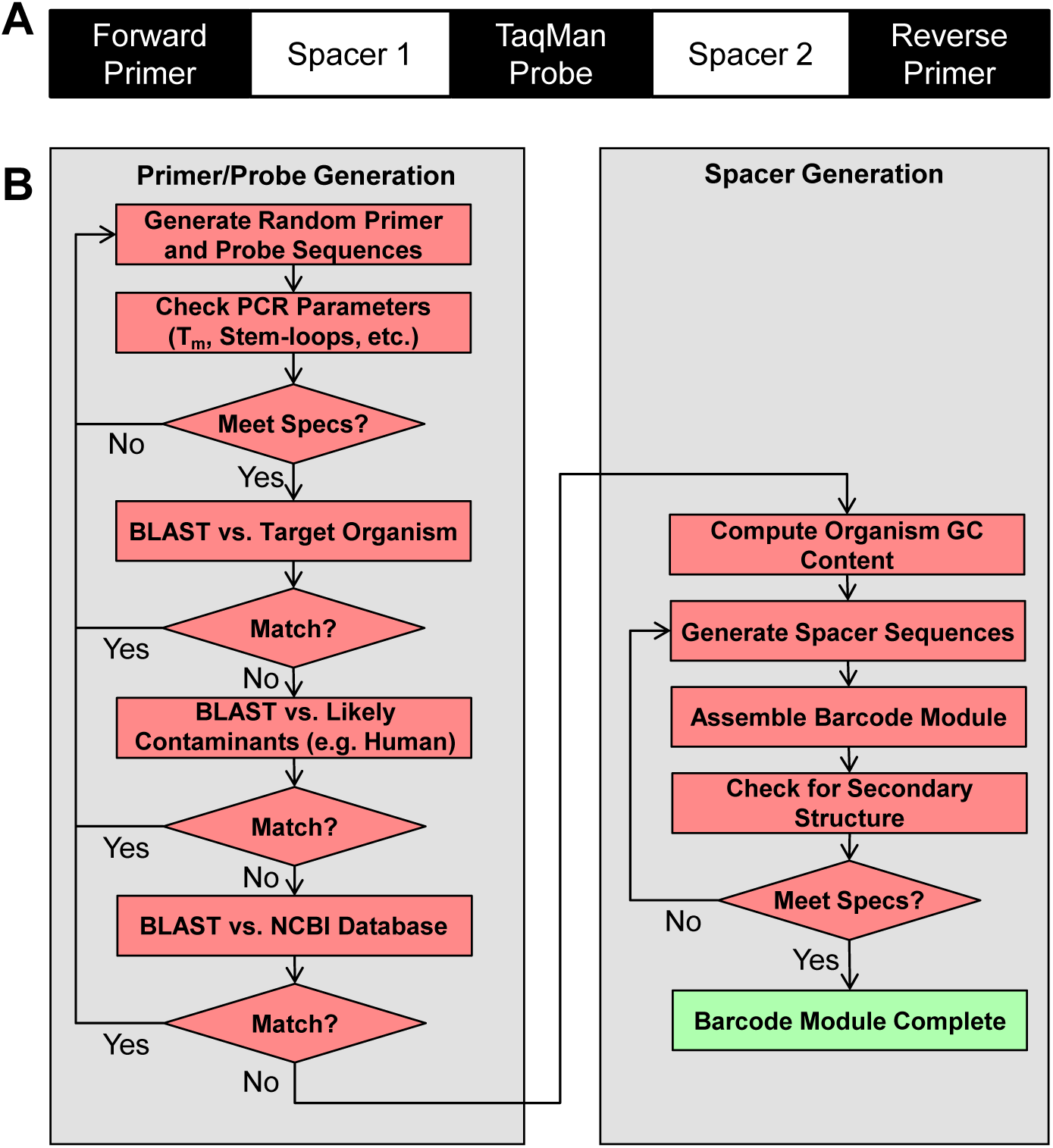
Overview of barcode design and algorithm work flow. A) Barcodes consist of two synthetic primer binding sites, a probe annealing site, and two spacer regions. Spacer regions can be adjusted to match the overall G+C content of the organism to be barcoded. B) barCoder algorithm work flow.

### Algorithm design

The barCoder algorithm workflow is depicted in Figure 1B. The algorithm starts by generating random primers to meet PCR-related specifications. Unlike typical primer design where primer sequences are constrained by an existing sequence of interest, here there is almost complete freedom to design primers that have ideal PCR properties. Using an approximate T_m_ formula (see Methods), the number of A’s+T’s and G’s+C’s needed to satisfy the specified T_m_ value can be calculated for primers within user-adjustable constraints on length and G+C content. From this set, a sequence meeting these constraints is randomly generated and screened for several PCR-related properties, such as maximum homopolymer repeats and secondary structure (Table 1). If any requirements are not met, the sequence is discarded and a new sequence generated.

**Table 1.**
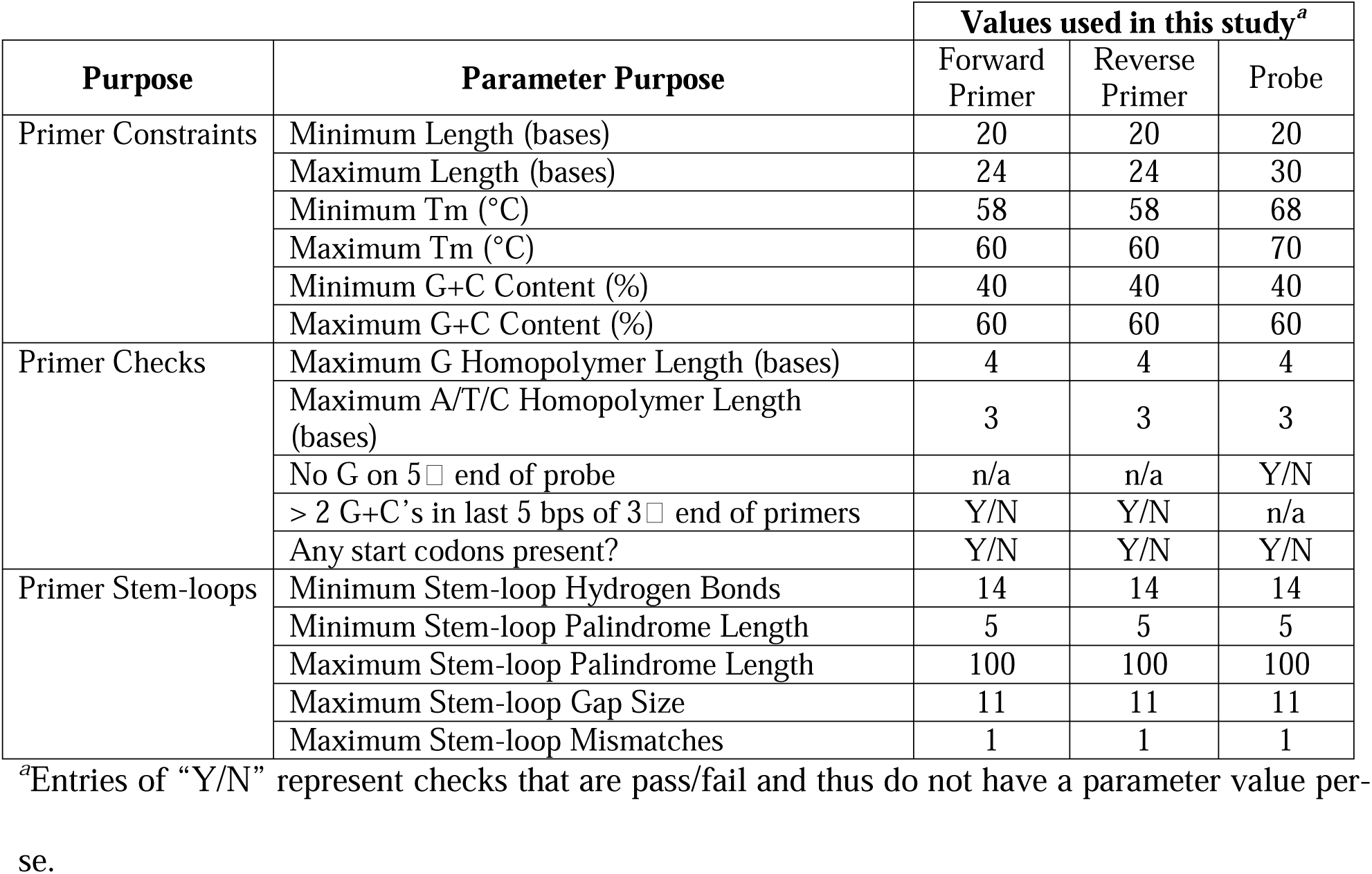
Algorithm parameters and default values for primer design.

A sequence that meets PCR restrictions is then tested for uniqueness. First, the sequence is compared to a list of other primers, which includes any primers already generated locally by the algorithm and an optional user-provided list of other primer sets of interest. Sequence “matches” are determined by comparing raw BLAST scores divided by the raw score of a perfect match to a user-adjustable threshold. If the sequence matches any existing primers above the threshold, the sequence is discarded and the process restarted. Second, the genome of the organism targeted for insertion is scanned for similar sequences by BLAST. Similarly, a set of additional genome sequences of organisms that may be likely to be present in a sample, such as common environmental background species or human, are scanned. Finally, the entire NCBI database is optionally scanned for similar sequences. A separate threshold for discarding a candidate sequence based on these BLAST results can be customized by the user, allowing more or less stringent criteria depending on project demands and acceptable CPU time in the case of very strict thresholds. Values used in this study are shown in Table 2.

**Table 2.**
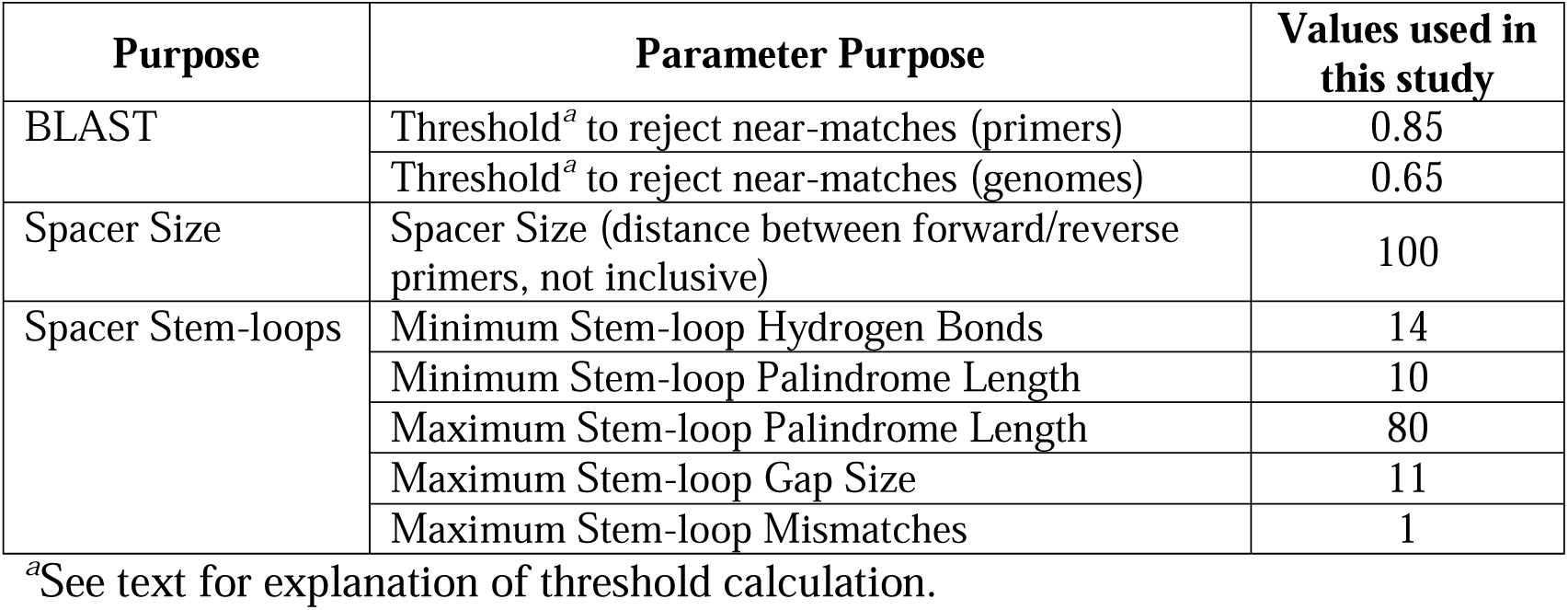
Algorithm parameters and default values other than for primer design.

| Purpose | Parameter Purpose | Values used in this study |
| --- | --- | --- |
| BLAST | Threshold<sup>a</sup> to reject near-matches (primers) | 0.85 |
|  | Threshold<sup>a</sup> to reject near-matches (genomes) | 0.65 |
| Spacer Size | Spacer Size (distance between forward/reverse primers, not inclusive) | 100 |
| Spacer Stem-loops | Minimum Stem-loop Hydrogen Bonds | 14 |
|  | Minimum Stem-loop Palindrome Length | 10 |
|  | Maximum Stem-loop Palindrome Length | 80 |
|  | Maximum Stem-loop Gap Size | 11 |
|  | Maximum Stem-loop Mismatches | 1 |

<sup>a</sup>See text for explanation of threshold calculation.

A sequence that meets all PCR and uniqueness requirements is accepted for use in the barcode. The algorithm cycles through this process to create each primer and the probe sequence, each with its own set of requirement parameters. Optionally, the forward primer can be set as constant for all barcodes in a given project. Once all three primer/probe sequences for a barcode have been generated, the spacer sequences are randomly generated such that the total length requirement is met and the G+C content of the full barcode matches the G+C content of the target organism. The final check scans for potential stem-loop structures in the barcode to limit challenges during genome insertion and during amplification of the sequence. Failing this check triggers regeneration of the spacer sequences.

### Experimental Validation

The barCoder algorithm was used to generate an initial set of 21 barcodes and corresponding qPCR detection primer/probe sets (sequences listed in Tables S1 and S2 in Additional file 1). Twelve of these barcodes were designed for *B. thuringiensis* serovar *kurstaki* (Btk), a surrogate for the biothreat agent *B. anthracis* with low-G+C content (35%, (33)). To demonstrate the utility of the barCoder algorithm to create barcodes for other organisms, including those with different G+C compositions, three barcodes each were designed for potential use in *Burkholderia pseudomallei* 1026b, (68% G+C content), *Yersinia pestis* CO92 (47%), and *Clostridium botulinum* Hall A (28%) (34-36).

Assay conditions for barcode Btk1 in the pIDTSMART-AMP plasmid backbone were optimized and subsequently standard curves were generated for all 21 TaqMan qPCR assays using the same conditions (Fig. 2 and Fig. S1 in Additional file 1). All of the assays of the barcodes in plasmids performed well with qPCR efficiencies ranging from 81.1% to 100.0%, strong linear relationships (R^2^ > 0.99), and no false positive results (Table 3). Limits of detection (LODs) were all below 50 copies (the lowest plasmid concentration tested), except for barcode Btk6, where the LOD was below 500 copies (Table 3). Select barcodes were also markerlessly incorporated into the chromosomes of potential target organisms: barcode Btk1 was integrated into both Btk and *B. anthracis* Sterne, and barcode Yp1 was inserted into a *pgm*^−^ derivative of *Y. pestis* CO92. Again, standard curves were generated for the TaqMan assays under the same conditions (Fig. 2). Assays using chromosomally-barcoded strains had efficiencies within the range observed for barcodes residing in plasmids (86.5% to 96.5%), R^2^ values above 0.99, and no false positives (Table 3). LODs were calculated as less than 15 copies and less than 2 copies for barcode Btk1 in the chromosomes of Btk and *B. anthracis* Sterne, respectively, and less than 25 copies for barcode Yp1 in the chromosome of *Y. pestis* CO92 *pgm*^−^ (Table 3). LODs are approximate as lower concentrations and Poisson distribution effects at low copy numbers were not thoroughly interrogated.

**Table 3.**
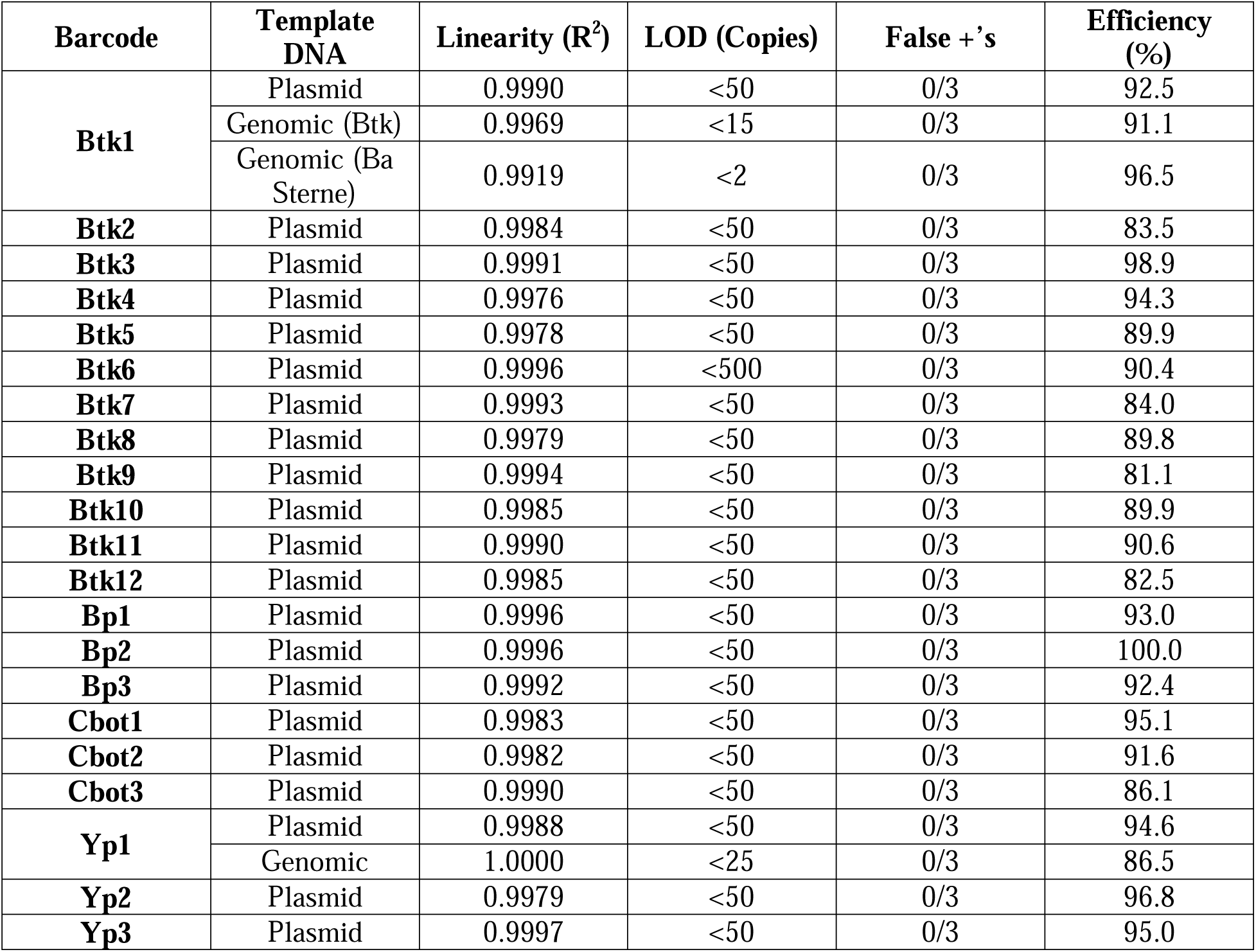
Evaluation of qPCR assays from generated standard curves.

**Figure 2.**
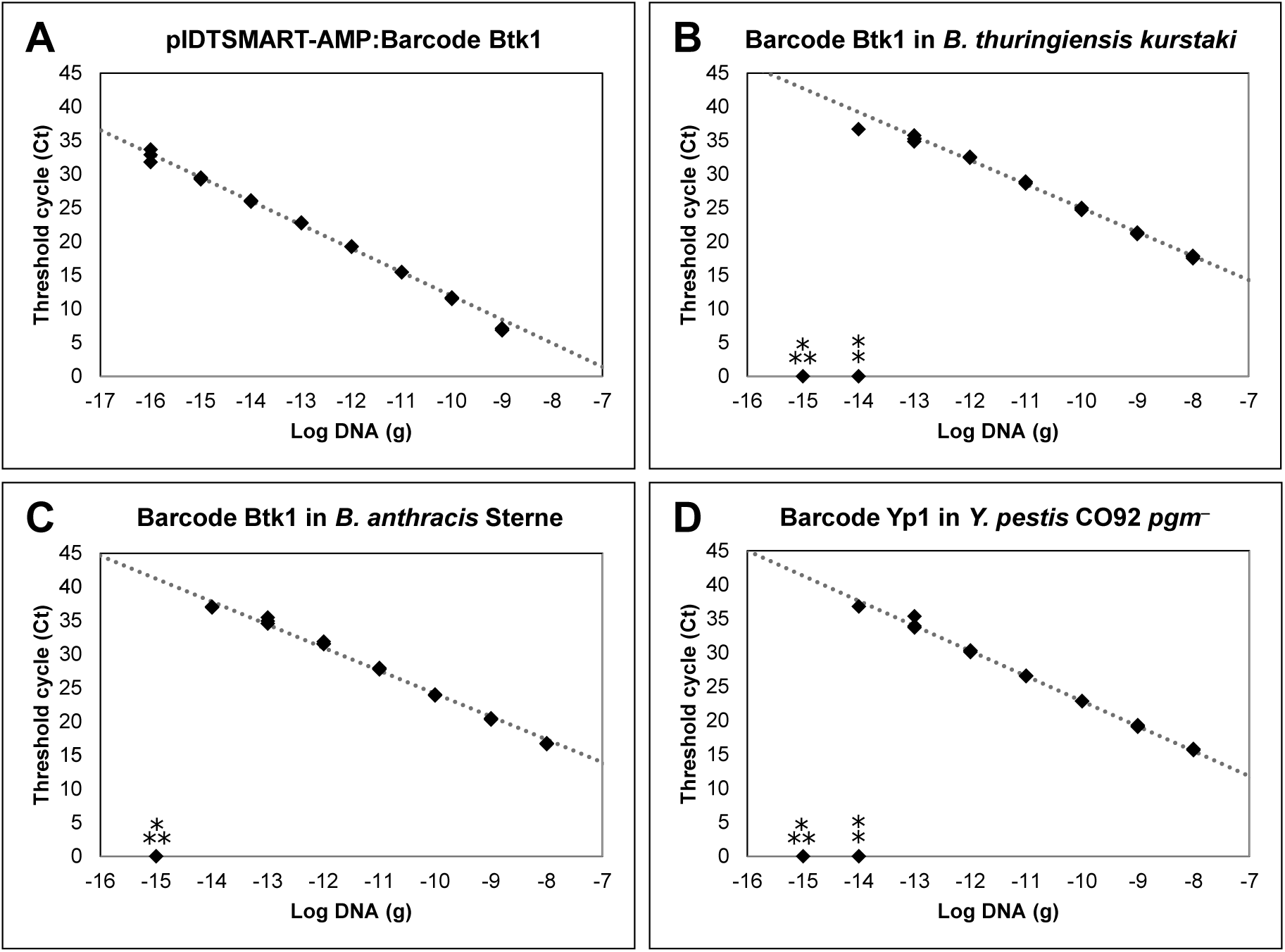
Representative qPCR assay standard curves. Curves were generated using A) barcode Btk1 in the pIDTSMART-AMP plasmid backbone, B) barcode Btk1 inserted into the *B. thuringiensis kurstaki* chromosome, C) barcode Btk1 inserted into the *B. anthracis* Sterne chromosome, and D) barcode Yp1 inserted into the *Y. pestis* CO92 *pgm*^−^ chromosome. For each standard curve, data from three replicates and a trendline are shown. Standard curves for the remaining barcodes in the plasmid backbone are shown in Figure S1 in Additional file 1. □ Ct value not determinable for 2/3 replicates. □ Ct value not determinable for 3/3 replicates.

To test the specificity of the TaqMan qPCR assays for the corresponding barcode, each of the 12 Btk assays were tested against all 12 Btk barcodes in plasmids. This cross-reactivity panel showed unique amplification of each Btk barcode with its cognate primer/probe set (Fig. 3). The TaqMan qPCR assay for barcode Btk1 was also tested against a panel of potential pathogens and environmental organisms (Table 4). Reactions containing the Btk strain with barcode Btk1 inserted in the chromosome, either alone or in the presence of an environmental matrix (DNA from a mock microbial community or DNA extracted from soil) showed robust positive results, while the Btk1 qPCR assay did not cross-react with any of the potential contaminants.

**Table 4.**
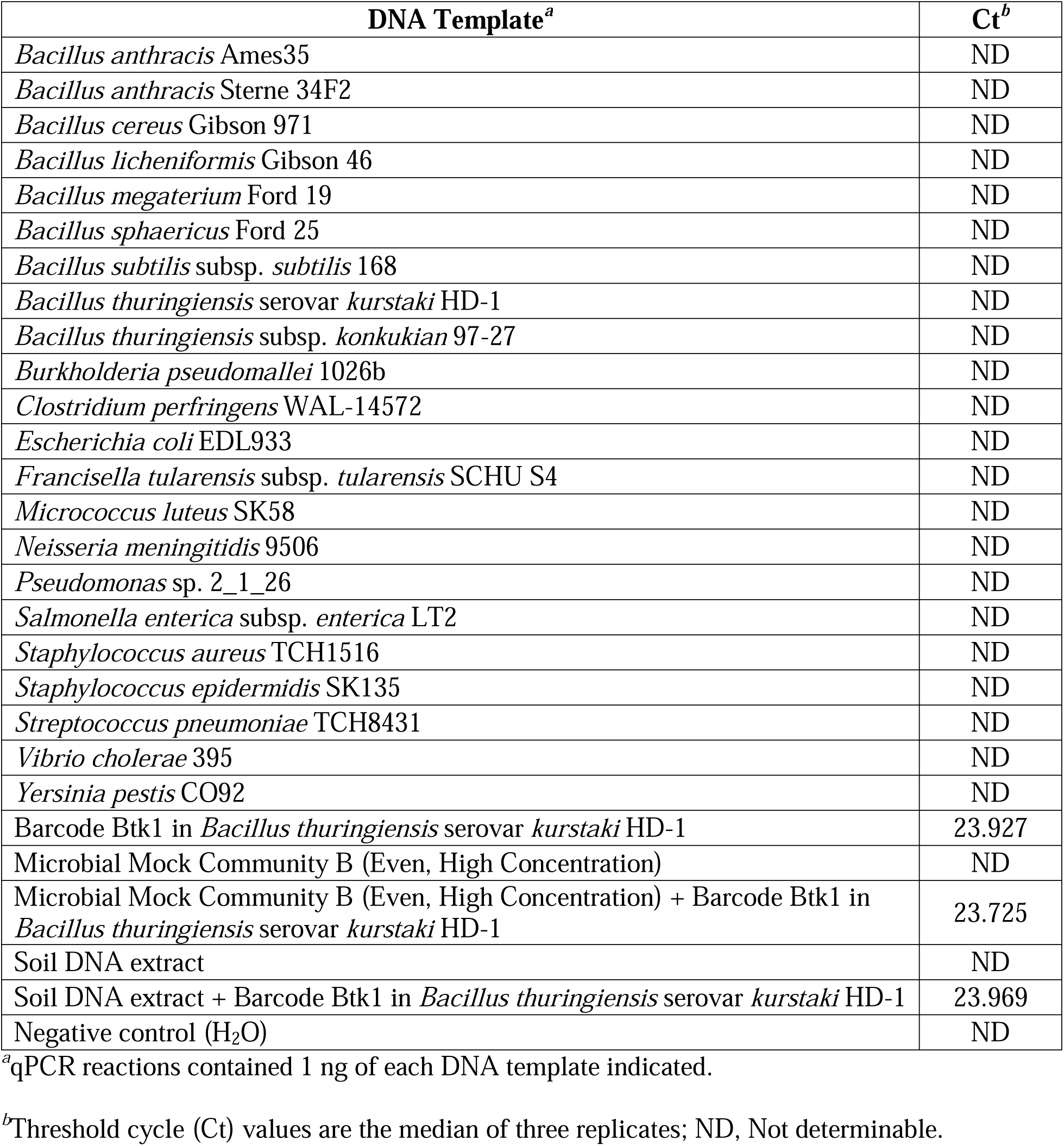
Cross-reactivity of the barcode Btk1 qPCR assay against a panel of potential pathogens and environmental contaminants.

| DNA Template<sup>a</sup> | Ct<sup>b</sup> |
| --- | --- |
| *Bacillus anthracis* Ames35 | ND |
| *Bacillus anthracis* Sterne 34F2 | ND |
| *Bacillus cereus* Gibson 971 | ND |
| *Bacillus licheniformis* Gibson 46 | ND |
| *Bacillus megaterium* Ford 19 | ND |
| *Bacillus sphaericus* Ford 25 | ND |
| *Bacillus subtilis* subsp. *subtilis* 168 | ND |
| *Bacillus thuringiensis* serovar *kurstaki* HD-1 | ND |
| *Bacillus thuringiensis* subsp. *konkukian* 97-27 | ND |
| *Burkholderia pseudomallei* 1026b | ND |
| *Clostridium perfringens* WAL-14572 | ND |
| *Escherichia coli* EDL933 | ND |
| *Francisella tularensis* subsp. *tularensis* SCHU S4 | ND |
| *Micrococcus luteus* SK58 | ND |
| *Neisseria meningitidis* 9506 | ND |
| *Pseudomonas* sp. 2_1_26 | ND |
| *Salmonella enterica* subsp. *enterica* LT2 | ND |
| *Staphylococcus aureus* TCH1516 | ND |
| *Staphylococcus epidermidis* SK135 | ND |
| *Streptococcus pneumoniae* TCH8431 | ND |
| *Vibrio cholerae* 395 | ND |
| *Yersinia pestis* CO92 | ND |
| Barcode Btk1 in *Bacillus thuringiensis* serovar *kurstaki* HD-1 | 23.927 |
| Microbial Mock Community B (Even, High Concentration) | ND |
| Microbial Mock Community B (Even, High Concentration) + Barcode Btk1 in *Bacillus thuringiensis* serovar *kurstaki* HD-1 | 23.725 |
| Soil DNA extract | ND |
| Soil DNA extract + Barcode Btk1 in *Bacillus thuringiensis* serovar *kurstaki* HD-1 | 23.969 |
| Negative control (H<sub>2</sub>O) | ND |

<sup>a</sup>qPCR reactions contained 1 ng of each DNA template indicated.
<sup>b</sup>Threshold cycle (Ct) values are the median of three replicates; ND, Not determinable.

**Figure 3.**
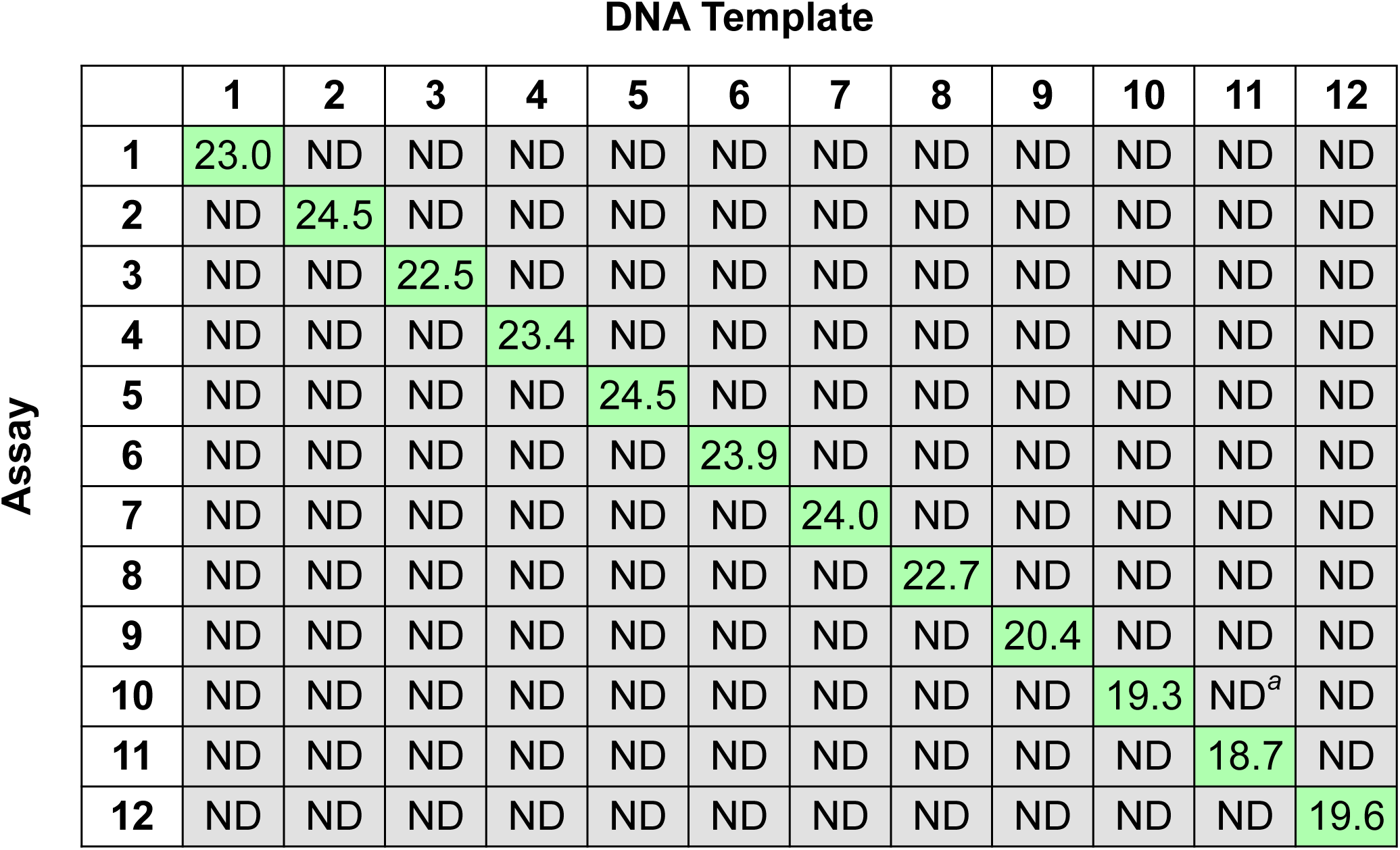
Cross-reactivity of the 12 Btk qPCR assays against the 12 Btk barcodes. For each qPCR reaction, 10^−12^g (∼450,000 copies) of the pIDTSMART-AMP plasmid backbone containing the Btk barcode indicated was used as DNA template. Threshold cycle (Ct) values shown are the median of three replicates; ND, Not determinable. ^*a*^1/3 replicates gave a Ct value of 37.087.

## Discussion

qPCR has become a standard technique for detection of microorganisms in the environment and for diagnosis of infection (37), and, as such, is an attractive detection technology that also allows a rapid evaluation of the relative abundance of a known microorganism within a sample. However, when conducting environmental fate studies, for example, these assays must discriminate from the endogenous or native microflora, which may be uncharacterized and present signatures similar to or cross-reactive with the signature selected for detection of the experimental strain. We sought in this report to utilize a bioinformatics strategy to generate specific amplicons that require minimal assay optimization and could be introduced into organisms with minimal to no cross-reactivity with environmental and microbial signatures.

Our approach to developing unique qPCR-compatible barcodes expanded upon our previous work, in which we appropriated synthetic signatures from published microarrays and developed PCR assays based on the unique sequences generated both by the tags themselves and by their insertion into the genome (24). Because those sequences were not designed *de novo* for use in PCR detection assays, we relied on the presence of a chromosomal primer binding site and the single synthetic sequence to generate suitable amplicons. As a result, considerable optimization of the assay conditions and primer sequences was necessary during the development of those strains, and the assay conditions for each tag required slightly different optimal conditions for detection. This situation was judged as suboptimal for the development of a more diverse panel of barcodes, as new assays would need to be developed for each new sequence.

We therefore sought to develop an algorithm that would enable the high-throughput generation of amplicon sequences that could use a single PCR assay condition, and in which relative proportion of each strain could be compared in a single test, e.g. across a single microplate. The assays would need to be specific to each barcode, and would need to be comparably sensitive with equivalent limits of detection. Using TaqMan™ qPCR chemistry and a stringent bioinformatic screening algorithm, we generated a panel of unique primer/probe combinations that exhibited the desired combination of selectivity, specificity, and sensitivity. Using conventional plasmids containing the barcodes as templates for the development of the assays, we demonstrate strong performance in linearity of response, sensitivity, and efficiency across 21 assays using conditions optimized for a single assay. No cross-reactivity was observed across a panel of 12 of these assays. We note that the odds of randomly generating a barcode that would react with a natural sequence is vanishingly small as three 20+ bp primer sequences would need to be closely matched in the correct orientation (>10^36^ possible sequences) with spacing appropriate for PCR amplification; nonetheless, sequences are screened for uniqueness to further minimize this possibility. Inserting two of the barcodes into three different genomes, we observed conserved performance compared to plasmid assays and LODs below 25 copies, which we believe to be conservative due to Poisson distribution effects at low copy numbers.

Our barcodes have a number of potential applications. Marking strains with unique artificial signatures could aid in protecting intellectual property, particularly for production strains whose development has required significant investment in metabolic and/or genetic optimization, perhaps in combination with other techniques such as DNA steganography (38). While not as information-rich as longer steganographic tags or watermarks (39, 40), qPCR barcodes have the advantage of not requiring further sequencing and informatic analysis to detect and/or verify their presence; they must simply be amplified using appropriate primers and probes. In one scenario, a set of barcodes could be inserted at defined intervals throughout a large DNA molecule used for information storage, and utilized to provide a preliminary indicator of the stability of the archive prior to full sequencing.

These sequences and their associated assays might also find use in forensic applications. In particular, one might imagine their use as molecular taggants that could be spiked into samples by field technicians, and their detection in DNA samples by the reference laboratories would serve to verify the origin of the sample. In a similar vein, these same tools could be used in the future for downstream attribution of accidental or deliberate release of organisms (41). Select agent strains, in particular, could be tagged, distributed to end-user communities, and then any material from the scene of a biocrime could be rapidly amplified using the library of primers and probes, enabling the rapid focus of investigative resources on those potential sources, while excluding the majority of the research laboratories that possess variants containing other barcodes. Any mechanism by which artificial genetic diversity can be introduced into the largely clonal populations of laboratory strains would be useful as all known acts of bioterrorism to date have utilized common laboratory strains (e.g. *B. anthracis* Ames Ancestor in the 2001 U.S. Mail/Amerithrax case; *S. enterica* serovar *typhimurium* 14028s in the Rajneeshi cult attacks of 1984) (42). In the case of the Amerithrax samples, discriminating between samples present in these laboratories relied on presence of several spontaneous mutants whose discovery and characterization required astute microbiologists and what were at the time Herculean sequencing efforts (43, 44). The deliberate incorporation of end-user-specific sequences into such commonly available strains could immeasurably speed identification of potential originating laboratories, would help investigators narrow their focus to a subset of potential sources, and would help exclude uninvolved laboratories working on similar research as potential sources. Furthermore, the presence of such signatures (and the knowledge that significant additional effort would be required to disguise the source of a sample) could deter potential malefactors within those laboratories even if the location, sequence, and properties of the sequence were known.

## Conclusions

To our knowledge, barCoder represents the first completely *in silico* method for generating both a synthetic target for qPCR and the primers/probe to amplify the target, and optimal assay conditions for detection of a diverse range of barcodes. We demonstrated that generated barcodes all perform well under a single set of assay conditions and show no cross-reactivity with themselves or environmental contaminants. Insertion of the barcodes into the genomes of three organisms of interest maintained the key properties of the barcodes. We anticipate barCoder finding utility in applications such as environmental fate studies, intellectual property, and microbial forensics.

## Methods

### Algorithm Design

All software was written in Perl. G+C and A+T constraints were calculated using the following formula:

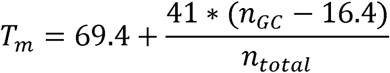

where *n*_*GC*_ is the number of G or C bases and *n*_*total*_ is the length of the primer.

Most bioinformatics functions were implemented using existing BioPerl modules. EMBOSS software, called by BioPerl, was used to predict stem-loop structures. All BLAST runs used default BioPerl parameters. Software is available on GitHub (https://github.com/ECBCgit/Barcoder).

### barCoder-designed elements and sources of DNA

All barcode modules were designed using the barCoder algorithm using the values listed in Tables 1 and 2. For the BLAST step against the target organism genome sequence, the following NCBI accession numbers were used: *B. thuringiensis* serovar *kurstaki*, NZ_CP010005.1; *B. pseudomallei* 1026b, NC_017831 (chromosome 1) and NC_017832 (chromosome 2); *C. botulinum* Hall A, NC_009495.1; *Y. pestis* CO92, NC_003143. All barcodes, primers, and probes were obtained from Integrated DNA Technologies, Inc. (IDT, Coralville, IA) and their sequences listed in Tables S1 and S2 in Additional file 1. Btk barcodes were designed with SacI/NheI restriction sites flanking the synthetic elements to facilitate later subcloning. Barcodes were received as “minigenes” inserted in the pIDTSMART-AMP plasmid backbone and were propagated in NEB® 5-alpha *E. coli* (New England Biolabs, Inc., Ipswich, MA) on Luria-Bertani (LB) agar and in LB broth with 100 µg/ml ampicillin at 37°C. Plasmid DNA was isolated using the QIAprep Spin Miniprep Kit (Qiagen, Hilden, Germany). DNA probes were ordered as PrimeTime double-quenched qPCR probes containing the 5□ FAM fluorophore, 3□ Iowa Black FQ quencher, and internal ZEN quencher. The sources of the DNA used for the cross-reactivity panel of pathogenic and environmental organisms are given in Table S3 in Additional file 1.

### qPCR

All qPCR experiments were run on an Applied Biosystems 7900HT Real-Time PCR System (Applied Biosystems, Foster City, CA) using Applied Biosystems MicroAmp optical 384-well reaction plates (catalog number 4309849) sealed with Applied Biosystems MicroAmp optical adhesive film (catalog number 4311971). Optimized 20 µL reactions included Applied Biosystems TaqMan® Universal PCR Master Mix (catalog number 4304437), forward and reverse primers each at a final concentration of 900 nM, DNA probe at a final concentration of 250 nM, 1 µL DNA template at the indicated concentration, and nuclease-free water. TaqMan assays used the following thermocycler protocol: 1 cycle of 50°C for 2 min, 1 cycle of 95°C for 10 min, and 40 cycles of 95°C for 15 sec and 55°C for 1 min. The standard curve properties of each assay were assessed by performing 10-fold serial dilutions of the template DNA in nuclease-free water. Efficiency and linearity (R^2^) values for each qPCR standard curve were calculated using the median Ct of three replicates for each template DNA dilution. Data points corresponding to the highest amount of template DNA tested (10^−8^ g for genomic DNA, 10^−9^ g for plasmid DNA) were omitted from these analyses in all cases as the Ct values tended to be non-linear with the other data points of the standard curve. LODs were conservatively estimated using the lowest amount of template DNA tested that produced a Ct value < 40 for all three replicates.

### Construction of genomically-barcoded strains

*B. thuringiensis* and *B. anthracis* strains were routinely cultured on Brain Heart Infusion (BHI) agar and in BHI broth at 30°C (*B. thuringiensis*) or 37°C (*B. anthracis*). Unless otherwise indicated, *Y. pestis* strains were grown on BHI agar or Tryptic Soy Agar (TSA), and in BHI broth at 28–30°C. Genomic DNA was extracted using the UltraClean Microbial DNA Isolation Kit (MOBIO Laboratories, Inc., Carlsbad, CA). Barcode Btk1 was selected to construct a strain in which the barcode was markerlessly incorporated into the chromosome of *B. thuringiensis* serovar *kurstaki* HD-1 (24, 33) (obtained from the DoD Unified Culture Collection (https://www.usamriid.army.mil/ucc/)). The insertion was generated at the same locus that was identified and modified in our previous report (within Target 1, (24)). This corresponds to an insertion between positions 4,834,064 and 4,834,065 of RefSeq accession number NZ_CP010005.1. Plasmid pRP1028-T1-PL (sequence provided in Additional file 2), a derivative of pRP1028 (45), was designed specifically for incorporating synthetic elements within this target region of the Btk chromosome and was synthesized by DNA2.0 (Menlo Park, CA). Plasmid pRP1028-T1-PL contains 1,550 bp of DNA homologous to the Btk chromosomal insertion region between the pRP1028 HindIII and BamHI sites, as well as a 36-bp polylinker within the homology region. Following digestion of plasmid pIDTSMART-AMP:Barcode Btk1 with SacI and NheI, the Btk1 barcode was gel extracted (QIAquick Gel Extraction Kit, Qiagen, Hilden, Germany) and ligated with pRP1028-T1-PL that had been digested with the same restriction enzymes. This pRP1028-T1-PL derivative containing barcode Btk1 was introduced into Btk, and the barcode was incorporated into the chromosome using the markerless allelic exchange strategy described previously (45). Successful barcode integration into the Btk chromosome was verified by PCR amplification of the target locus and SacI/NheI digestion of the resulting amplicon. Construction of a strain of *B. anthracis* Sterne 34F2 with barcode Btk1 in the chromosome was previously published (46).

For construction of a strain of *Y. pestis* CO92 *pgm*^−^ with barcode Yp1 markerlessly inserted in the chromosome, the locus between the convergently transcribed genes YPO0388 and YPO0392a (RefSeq accession number NC_003143.1) was selected using rules adopted from Buckley et al. (24), in combination with the PATRIC database (47) and available transcriptome sequencing (RNA-seq) data (SRA accession numbers SRR1013703, SRR1013704, SRR1013705, SRR1041589), to identify a potentially neutral insertion region. The barcode was then inserted into the chromosome between positions 406,742 and 406,743 via the method described by Sun et al. (48), which utilizes λ Red recombination and *sacB* counterselection. Briefly, plasmid pKD46 (CGSC #7739, (48)) containing the genes for λ Red recombination was electroporated into a strain of *Y. pestis* CO92 *pgm*^−^ (strain R88, Robert Perry, University of Kentucky). A linear DNA fragment containing a *cat*-*sacB* cassette flanked by DNA homologous to the *Y. pestis* chromosomal insertion region was electroporated into this pKD46-containing strain of *Y. pestis*, and successful integrants were selected on media containing chloramphenicol. Following electroporation with a linear DNA fragment containing barcode Yp1 flanked by homologous DNA and selection on media containing sucrose, the *cat*-*sacB* cassette in the chromosome was replaced with the barcode. The resulting strain was subsequently cured of pKD46, and successful barcode insertion was verified by PCR amplification and sequencing. Whole-genome sequencing (MiSeq, Illumina) was also performed to confirm the absence of off-target modifications. Primers used to construct this barcoded strain of *Y. pestis* are listed in Table S4 in Additional file 1. To generate the *cat*-*sacB* cassette, the *cat* gene was PCR amplified from plasmid pKD3 (CGSC #7631, (49)) with primers #1 and #2, and the *sacB* gene was PCR amplified from plasmid p88171 (synthesized plasmid with pJ207 backbone and *sacB* gene from *Bacillus subtilis*, DNA 2.0, Menlo Park, CA) with primers #3 and #4. The two PCR amplicons were purified (QIAquick PCR Purification Kit, Qiagen, Hilden, Germany) and joined together by overlap extension PCR (50) using primers #1 and #4. The *cat*-*sacB* cassette was gel extracted and cloned between the SacI and BamHI sites of pUC19 to create plasmid pCBV4. To construct the *cat*-*sacB* cassette flanked by homologous DNA, the *cat*-*sacB* cassette was PCR amplified from pCBV4 with primers #5 and #6. Approximately 500 bp flanking each side of the barcode insertion point were separately PCR amplified from the *Y. pestis* CO92 *pgm*^−^ chromosome; primers #7 and #8 were used to amplify upstream DNA, and primers #9 and #10 were used to amplify downstream DNA. The three purified PCR amplicons (up flanking region, *cat*-*sacB* cassette, and down flanking region) were joined together by overlap extension PCR (50) using primers #7 and #10, and the resulting amplicon was gel extracted and cloned into the pCR™4Blunt-TOPO® vector (Invitrogen, Carlsbad, CA) to generate plasmid pCBV6. The linear DNA fragment containing the *cat*-*sacB* cassette flanked on both sides by *Y. pestis* CO92 *pgm*^−^ DNA was PCR amplified from pCBV6 with primers #7 and #10. To create barcode Yp1 flanked by homologous DNA, the barcode was PCR amplified from the synthesized plasmid pIDTSMART-AMP:Barcode Yp1 using primers #11 and #12. Approximately 500 bp flanking each side of the barcode insertion point were separately PCR amplified from the *Y. pestis* CO92 *pgm*^−^ chromosome; primers #7 and #13 were used to amplify upstream DNA, and primers #10 and #14 were used to amplify downstream DNA. The three purified PCR amplicons (up flanking region, barcode, and down flanking region) were joined by overlap extension PCR (50) using primers #7 and #10, and the resulting amplicon was gel extracted and cloned into the pCR™4Blunt-TOPO® vector (Invitrogen, Carlsbad, CA) to generate plasmid pCBV9. The linear DNA fragment containing barcode Yp1 flanked on both sides by *Y. pestis* CO92 *pgm*^−^ DNA was PCR amplified from pCBV9 with primers #7 and #10.

## Declarations

### Ethics approval and consent to participate

Not applicable

### Consent for publication

Not applicable

## Availability of data and material

All raw data to the level of Ct values that were generated and analyzed during this study are included in the article and its additional files.

## Authors’ contributions

CBB inserted barcodes into *B. anthracis* and *Y. pestis*, designed and performed qPCR experiments, analyzed data, and wrote the manuscript. MWL developed the algorithm, designed qPCR experiments, analyzed data, and wrote the manuscript. SEK optimized qPCR protocols. TDPG inserted barcode Btk1 into Btk. ATL updated the algorithm for publication. HSG conceived and led the project, analyzed data, and wrote the manuscript.

## Acknowledgements

We thank F. Chris Minion (Iowa State University) for the generous gift of *Y. pestis* strain R88. We also thank Mark Karavis (CCDC CBC) for whole-genome sequencing, and Pierce Roth, Edward Fochler (DCS Corp/CCDC CBC), and Michael Krepps (BrightEdge Investments) for bioinformatics support.

## Additional Files

Additional file 1.docx: Supplementary Tables and Figure. Tables S1 contains sequences for the 21 barcodes designed with the barCoder algorithm. Table S2 contains primer and probe sequences for each barcode module. Table S3 provides the sources of DNA used in the cross-reactivity panel shown in Table 4. Table S4 contains sequences of the primers used to construct the barcoded strain of *Y. pestis* CO92 *pgm*^−^. Figure S1 shows additional qPCR standard curves.

Additional file 2.gbk: Plasmid pRP1028-T1-PL sequence. This file contains the complete annotated sequence for the plasmid designed for incorporating synthetic elements within the target region of the Btk chromosome.

Additional file 3.xlsx: Standard curve data. This file contains Ct values and calculations used to generate and analyze qPCR standard curves for all 21 barcodes (in the plasmid backbone and inserted in the genome, if applicable).

Additional file 4.xlsx: Raw data for the cross-reactivity panel of Btk barcodes and qPCR assays. This file contains raw Ct values from the cross-reactivity panel of the 12 Btk qPCR assays against the 12 Btk barcodes.

Additional file 5.xlsx: Raw data for the Btk1 qPCR assay cross-reactivity panel. This file contains the raw Ct values from the cross-reactivity panel of the barcode Btk1 qPCR assay against a panel of potential pathogens and environmental contaminants.

