## Additional File 1 for "barCoder: a tool to generate unique, orthogonal genetic tags for qPCR detection"

^†^Corresponding author

Table S1. Barcode sequences.

| **Barcode** | **Sequence (5ʹ to 3ʹ)***^a^* |
| --- | --- |
| Btk1 | GAGCTCTGTCTCCAACTCCCAGTATCTTATTTTCTTAATAGATTATATCAAGACTATTAATTTTACCCGACGCAGGACCCCACTAAGTGATTTCAATATTTAGAAATTATTGTAAAATATCTTAACTTCGCTGCCTCAAACC**GCTAGC** |
| Btk2 | GAGCTCTCACACACCTATTTACTCCCTATTTTCAATTGTGTTTTAATTTAAGAGGATATATTACGCTCAAGGAAGCCTTCCCAAGTTCACACACTTATCTGTATATATAAATTTAAGAAAACAAAGGCGTGTACTAGACTTGCTT**GCTAGC** |
| Btk3 | GAGCTCAAACAGACGAGTCCTGAGAATCAATTATTTATCTAACTACAAACTATATTACAGTATTTACAACAACCAGAAAGACCCCTCCGGGAAATAATATTGTAAAATTAAAAGGTCTGAACAATATAGCGGAAGAGCAGTCTCTTAT**GCTAGC** |
| Btk4 | GAGCTCGTTGGTAGGCCAAACTACTTCTTACTTATAAGTTCGAAGATTTTAATATTGAATATCGATACAGGATCACGAGCGCCTCTAACTATAAACAATATACCTATTTCCAAAAGAAATAAAAATCGACAGTTATAGGAACTAGAC**GCTAGC** |
| Btk5 | GAGCTCCGATTTCTCAACCTATACACTGAGTGAGTTGTATTGTTAAAAAATCTCTTAAATAAACTCCGAGGTGATCACCAGAGTTCAGACTCAAGCTAATAAATCAACTTATATAAAATTAAGTCAGAAGTTCGGGTAATCAGAAGTT**GCTAGC** |
| Btk6 | GAGCTCGTCCGCCGCCCAATTTATATAAGTTTAAAAAAACTTAATAAAGATATTGTTCTAGGCCGCTTAAGCGCGCAGTTCTATTACTTTTTCTTTTTAAAATCAATATAATTTAGCCTAAACGCCGCTCAACGATTCTTTA**GCTAGC** |
| Btk7 | GAGCTCTGAGTGACAGGTGATCGACTTTATATTTATTATATTAGTAAGTCTTATCTACTATTAGCCTTCTAGTGAGTCGCCAACCGAGTTTTAACTGATTACTACTATAAATTTCAATTAATTTTTGGAGTAGTGCGCCGTTT**GCTAGC** |
| Btk8 | GAGCTCATTCTCACGGCTTCGACTTTCGTAAAAATAGTTTAGCTAAACTGAATTATTTTTAATTCGGTCCCCACTAATACCCCAGAGTGAGAAAGTTACACAATTTGAATTATATTTTAGTAATTTGCCTTAACTGACGGAACTCA**GCTAGC** |
| Btk9 | GAGCTCTCTGCCTGGAGTGATTTAGAAATATAGTTACAAACTATTTTAGAATATAATAGTGAGTACGTCGCTATATATCTCCCCACCCACCTATAAAGAATCCTAACAGTTTCTAGAAATAAAAATTAGTCTAATTCGTAGGCAACAG**GCTAGC** |
| Btk10 | GAGCTCTAATATCGGCAGACGTGCTGTTTACTGAAATTAAACTTATTAGAACTAAATACTACTTACCGTTAAGCTACAGGACACCGCAACGAAAAATCACGATAATCTATATAATTACCGTATAAAAACTGAATCAAACTGGGTGATCT**GCTAGC** |
| Btk11 | GAGCTCACGGAGACGGTCTGTTATTGTAATCAAAAGACTAAACAAAAATTTCTAAAATTAAAGCGGCACTGCGCTAAACTCCCTTAGATATATTTTGCTTATAATAAAATACTAAGTACAATCTCGGCTATCCCTATTCAGT**GCTAGC** |
| Btk12 | GAGCTCCGTCGGTATTGCACCTTACAACGATTAATTTATAATAATTAAGTTGTATTATCTCTGAAGCGTCCTCTAGTCCACTTGGAGCAAAATTATTATCACTTATCTAATCTTAAATTTTACACGTGGATCACAACGCTAT**GCTAGC** |
| Bp1 | GCCTCACTTGAGGTAATTAGAATCTGCGGGGCGCGCGGGCGAACCCGACGGCGGGGGATCGCGACTCACTCCTACACGTAAAAGGGCTGCCCGTCGCTCCCGCCCGGGGGGGCTGCCGGGGATTGCCCCTTCGTTAAACCA |
| Bp2 | ATATAGCCGCGGACAGATCTTACGGCCGAGTCCGCCCCGGCAGCGCCCCCGCCTAGGGAGTCCGACTTTCAAATCACCGCACCCAGCCCGGCCGCCGCTAGCTCGGGAGGGCGGTGTACCAAGCCCGGTTGTTTCA |
| Bp3 | AGGCAGGTGGCGAGTAAATAGGTCTCGCTCGTCGCTGGGACCGCCCCGGGTGGGCTGCCGATAGGCCCTTACAACTCTGCGGCCGCCCGTGCTGGCTGGCAGCAGTCGACCCGGCCCTGGACACGCTCAAGGTTAT |
| Cbot1 | GTGGTGCGAGACTGTAAAGATAATTAAAAATATTAATTTAGTTTATTTTTAAATAAGGGTCACACGTTGCCCTTCTTCTATCGTAATTAAATTTAAATTAATAATAAATTTTAAGTAACGACGTTGTTTAGGGTTACG |
| Cbot2 | TTCGCTCTGCAAATTTGAAATCGTTATTAGATTAAAATTACTTTATTAAAAAAATCTTATTGCCCAAGGATACGTGCACCTCTAGGAAAATTTTTAAGATTATATTTTAATAGTATTTTAGAAAGCAGACGAAAGATTGACTAC |
| Cbot3 | CGGAATAGTGAACGGTTCCTTTTAATTATTTAAATTATAAAAAAAATAATCAATATTGAAGCGTCGTCCTAGCCCTTATTGCAGATTTCTTTAAAATAAAATAATTATATAATTATAAAAGGAGTCAGTAGCTGTTGTGA |
| Yp1 | AAGATTACGAGTTGGCACGAAATTCGAGTATTATAGGTCTAGCTCTGCGAGAAGACCGCTTAAGCTCGGGTACCTATCCACCGAGTACCCGCGCTCTTAGCTAGATTAGTTCTTACACCCTTCGTAAAGTTACCGGTTGAAGA |
| Yp2 | TGTTGTCGGAGACCTATTCTGTTTTCTGTGAGGGACTAAGAATAGCCTTACTCCTGCCGTACCCGAAGTGACGTACTTCTCCTGAGCCTCCGAGTATTATTGGATCACCGCAACTGTTTGGACAGAGACTGGTTAATTATTG |
| Yp3 | CCGTAGCTGTTGATCGTCAAAAATCACGATTCTTTGGTACGAGGTCAATCACCACACCAGGTGGCCCAAATCTTATACGGAACCTCGGCTTTGTAGAACGATTCCTAACGTGAAGTCAGGCAACTTGCTAAGAAGTCGAA |

*^a^*SacI restriction sites are underlined; NheI restriction sites are shown in bold.

Table S2. Barcode primer and probe sequences.

| **Barcode** | **Forward Primer (5ʹ to 3ʹ)** |
| --- | --- |
| Btk1 | TGTCTCCAACTCCCAGTATC |
| Btk2 | TCACACACCTATTTACTCCCTAT |
| Btk3 | AAACAGACGAGTCCTGAGAATC |
| Btk4 | GTTGGTAGGCCAAACTACTTC |
| Btk5 | CGATTTCTCAACCTATACACTGA |
| Btk6 | GTCCGCCGCCCAATTTATA |
| Btk7 | TGAGTGACAGGTGATCGACTT |
| Btk8 | ATTCTCACGGCTTCGACTTTC |
| Btk9 | TCTGCCTGGAGTGATTTAGAAAT |
| Btk10 | TAATATCGGCAGACGTGCTGTT |
| Btk11 | ACGGAGACGGTCTGTTATTG |
| Btk12 | CGTCGGTATTGCACCTTAC |
| Bp1 | GCCTCACTTGAGGTAATTAGAAT |
| Bp2 | ATATAGCCGCGGACAGATC |
| Bp3 | AGGCAGGTGGCGAGTAAATA |
| Cbot1 | GTGGTGCGAGACTGTAAAGAT |
| Cbot2 | TTCGCTCTGCAAATTTGAAATCG |
| Cbot3 | CGGAATAGTGAACGGTTCCTTT |
| Yp1 | AAGATTACGAGTTGGCACGAAAT |
| Yp2 | TGTTGTCGGAGACCTATTCTGT |
| Yp3 | CCGTAGCTGTTGATCGTCAAA |
|  | **Reverse Primer (5ʹ to 3ʹ)** |
| Btk1 | GGTTTGAGGCAGCGAAGTTA |
| Btk2 | AAGCAAGTCTAGTACACGCC |
| Btk3 | ATAAGAGACTGCTCTTCCGCTA |
| Btk4 | GTCTAGTTCCTATAACTGTCGAT |
| Btk5 | AACTTCTGATTACCCGAACTTCT |
| Btk6 | TAAAGAATCGTTGAGCGGCGT |
| Btk7 | AAACGGCGCACTACTCCAAA |
| Btk8 | TGAGTTCCGTCAGTTAAGGCA |
| Btk9 | CTGTTGCCTACGAATTAGACTAA |
| Btk10 | AGATCACCCAGTTTGATTCAGTT |
| Btk11 | ACTGAATAGGGATAGCCGAG |
| Btk12 | ATAGCGTTGTGATCCACGTG |
| Bp1 | TGGTTTAACGAAGGGGCAATC |
| Bp2 | TGAAACAACCGGGCTTGGTA |
| Bp3 | ATAACCTTGAGCGTGTCCAG |
| Cbot1 | CGTAACCCTAAACAACGTCG |
| Cbot2 | GTAGTCAATCTTTCGTCTGCTTT |
| Cbot3 | TCACAACAGCTACTGACTCCTT |
| Yp1 | TCTTCAACCGGTAACTTTACGAA |
| Yp2 | CAATAATTAACCAGTCTCTGTCC |
| Yp3 | TTCGACTTCTTAGCAAGTTGCC |
|  | **Probe (5ʹ to 3ʹ)** |
| Btk1 | TTTACCCGACGCAGGACCCCACTAAG |
| Btk2 | CGCTCAAGGAAGCCTTCCCAAGTTCACA |
| Btk3 | TTACAACAACCAGAAAGACCCCTCCGGG |
| Btk4 | CGATACAGGATCACGAGCGCCTCTAAC |
| Btk5 | CTCCGAGGTGATCACCAGAGTTCAGACT |
| Btk6 | AGGCCGCTTAAGCGCGCAGTTCTATTAC |
| Btk7 | TAGCCTTCTAGTGAGTCGCCAACCGAGT |
| Btk8 | TTCGGTCCCCACTAATACCCCAGAGTGA |
| Btk9 | TACGTCGCTATATATCTCCCCACCCACC |
| Btk10 | TACCGTTAAGCTACAGGACACCGCAACG |
| Btk11 | AGCGGCACTGCGCTAAACTCCCTTAG |
| Btk12 | CTGAAGCGTCCTCTAGTCCACTTGGAG |
| Bp1 | ATCGCGACTCACTCCTACACGTAAAAGGG |
| Bp2 | TAGGGAGTCCGACTTTCAAATCACCGCAC |
| Bp3 | TGCCGATAGGCCCTTACAACTCTGCG |
| Cbot1 | AGGGTCACACGTTGCCCTTCTTCTATCGT |
| Cbot2 | ATTGCCCAAGGATACGTGCACCTCTAGG |
| Cbot3 | TGAAGCGTCGTCCTAGCCCTTATTGCAG |
| Yp1 | CTTAAGCTCGGGTACCTATCCACCGAG |
| Yp2 | CGTACCCGAAGTGACGTACTTCTCCTG |
| Yp3 | ACCAGGTGGCCCAAATCTTATACGGAACC |

Table S3. Sources of DNA for cross-reactivity panel.

| **DNA Template** | **Source** | **Catalog #** |
| --- | --- | --- |
| *Bacillus anthracis* Ames35 | BEI*^a^* | NR-10450 |
| *Bacillus anthracis* Sterne 34F2 | S. Leppla (NIH)*^b^* | − |
| *Bacillus cereus* Gibson 971 | BEI*^a^* | NR-4198 |
| *Bacillus licheniformis* Gibson 46 | BEI*^a^* | NR-2546 |
| *Bacillus megaterium* Ford 19 | BEI*^a^* | NR-2543 |
| *Bacillus sphaericus* Ford 25 | BEI*^a^* | NR-2548 |
| *Bacillus subtilis* subsp. *subtilis* 168 | BGSC*^b^* | 1A1 |
| *Bacillus thuringiensis* serovar *kurstaki* HD-1 | BEI*^b^* | NR-610 |
| *Bacillus thuringiensis* subsp. *konkukian* 97-27 | BEI*^a^* | NR-12315 |
| *Burkholderia pseudomallei* 1026b | BEI*^a^* | NR-9321 |
| *Clostridium perfringens* WAL-14572 | BEI*^a^* | HM-310D |
| *Escherichia coli* EDL933 | BEI*^a^* | NR-2648 |
| *Francisella tularensis* subsp. *tularensis* SCHU S4 | BEI*^a^* | NR-3015 |
| Microbial Mock Community B (Even, High Concentration) | BEI*^a^* | HM-276D |
| *Micrococcus luteus* SK58 | BEI*^a^* | HM-114D |
| *Neisseria meningitidis* 9506 | BEI*^a^* | NR-48805 |
| *Pseudomonas* sp. 2_1_26 | BEI*^a^* | HM-214D |
| *Salmonella enterica* subsp. *enterica* LT2 | BEI*^a^* | NR-4218 |
| Soil DNA extract | Abingdon, MD*^c^* | − |
| *Staphylococcus aureus* TCH1516 | BEI*^a^* | NR-12252 |
| *Staphylococcus epidermidis* SK135 | BEI*^a^* | HM-118D |
| *Streptococcus pneumoniae* TCH8431 | BEI*^a^* | HM-145D |
| *Vibrio cholerae* 395 | BEI*^a^* | NR-15694 |
| *Yersinia pestis* CO92 | BEI*^a^* | NR-2717 |

*^a^*Genomic DNA was obtained from the indicated commercial source; BEI, BEI Resources (Manassas, VA).

*^b^*Organism was obtained from the indicated source and genomic DNA extracted in-house using UltraClean Microbial DNA Isolation Kit (MOBIO Laboratories, Inc., Carlsbad, CA); BGSC, Bacillus Genetic Stock Center (Columbus, OH).

*^c^*Sample was collected at the indicated location and genomic DNA extracted in-house using DNeasy PowerSoil Kit (Qiagen, Hilden, Germany).

Table S4. Primers for constructing barcoded *Y. pestis* strain.

| **Primer #** | **Primer Name** | **Sequence (5ʹ→3ʹ)***^a^* |
| --- | --- | --- |
| 1 | pKD3_Cm-F | CTA**GAGCTC**TGTGACGGAAGATCAC |
| 2 | pKD3_Cm_SacB-R | CTATTAGACTCGAATAGGAACTTCGGAATAGG |
| 3 | p88171_SacB_Cm-F | CGAAGTTCCTATTCGAGTCTAATAGAATGAGGTCG |
| 4 | p88171_SacB-R | CAT**GGATCC**TGCCAATAGGATATCGGC |
| 5 | Cm_Yp2-F | *CAATCTACATCAACTTAC*CTGTGACGGAAGATCAC |
| 6 | SacB_Yp2-R | *GATGTACCTACATCA*TGCCAATAGGATATCGGC |
| 7 | Yp2_UpFlank_F | *GGATAGGGGATATGCTGC* |
| 8 | Yp2_UpFlank_Cm-R | GTCACAG*GTAAGTTGATGTAGATTGAATCAG* |
| 9 | Yp2_DownFlank_SacB-F | CCTATTGGCA*TGATGTAGGTACATCTGCGG* |
| 10 | Yp2_DownFlank_R | *CGTGAACTCTGCCGCAAG* |
| 11 | Yp2_Barcode1_F | *ACATCAACTTAC*AAGATTACGAGTTGGCAC |
| 12 | Yp2_Barcode1_R | *CCTACATCA*TCTTCAACCGGTAACTTTAC |
| 13 | Yp2_UpFlank_R | CTCGTAATCTT*GTAAGTTGATGTAGATTGAATCAG* |
| 14 | Yp2_DownFlank_F | CGGTTGAAGA*TGATGTAGGTACATCTGCGG* |

*^a^*Restriction sites are in bold. Bases in italics represent regions that align with the *Y. pestis* CO92 *pgm*^−^ chromosome. Underlined sequences align with barcode Yp1.

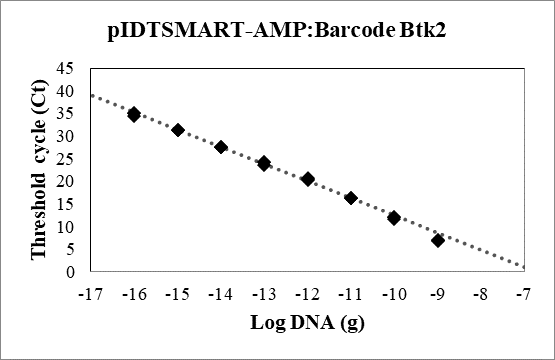

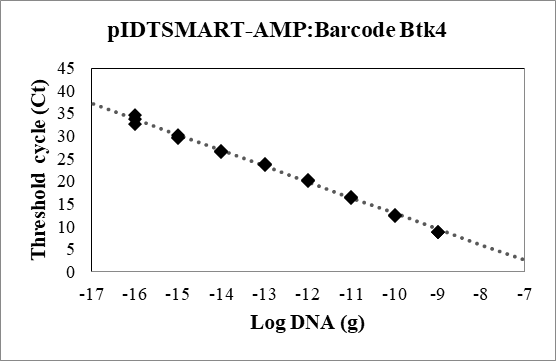

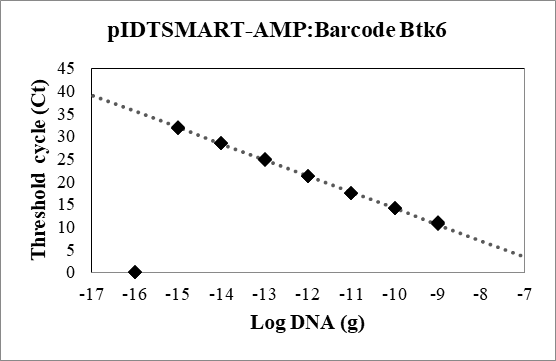

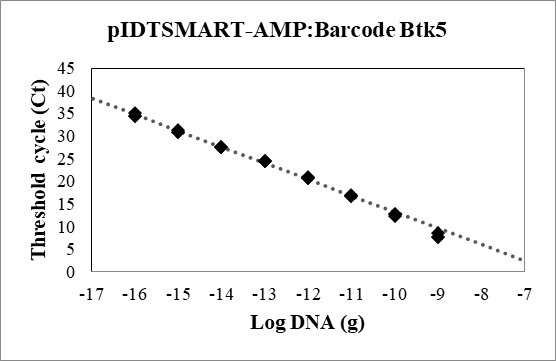

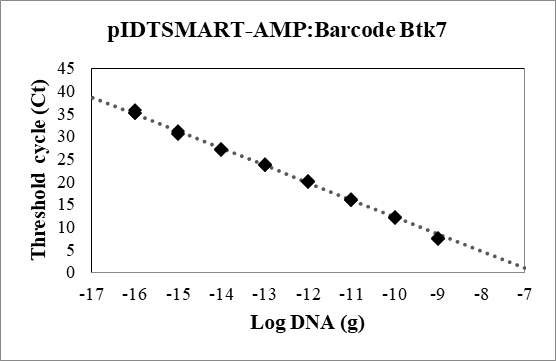

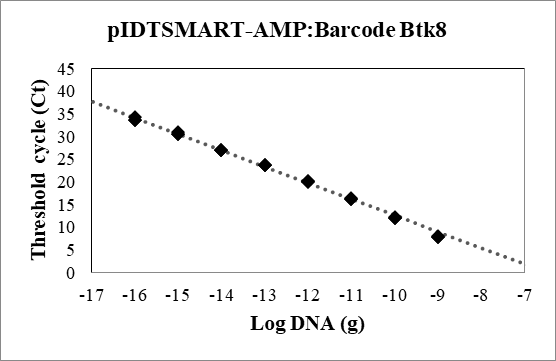

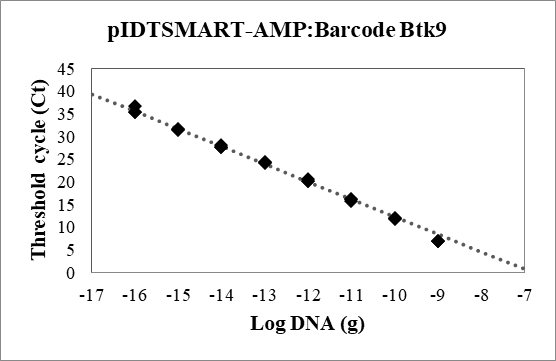

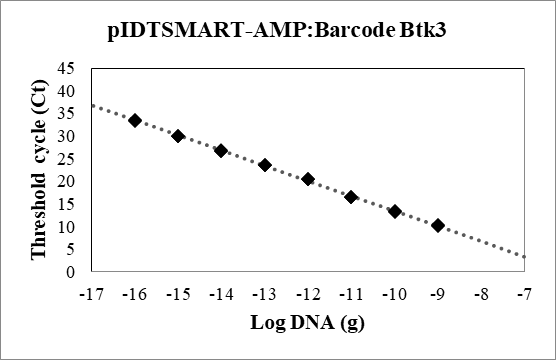

pIDTSMART-AMP:Barcode Btk2

Threshold cycle (Ct)

| Log DNA (g) | Ct |
| --- | --- |
| -9 | 6.91883 |
| -9 | 7.03757 |
| -9 | 7.07207 |
| -10 | 12.0964 |
| -10 | 11.7517 |
| -10 | 12.0066 |
| -11 | 16.2188 |
| -11 | 16.3239 |
| -11 | 16.223 |
| -12 | 20.3804 |
| -12 | 20.563 |
| -12 | 20.7023 |
| -13 | 24.2842 |
| -13 | 23.6747 |
| -13 | 24.1554 |
| -14 | 27.6013 |
| -14 | 27.5597 |
| -14 | 27.4945 |
| -15 | 31.3646 |
| -15 | 31.2726 |
| -15 | 31.2838 |
| -16 | 35.035 |
| -16 | 35.1662 |
| -16 | 34.4361 |

| Log DNA (g) | Trend (adj) |
| --- | --- |
| -18 | 42.7973 |
| -17 | 39.0043 |
| -16 | 35.2113 |
| -15 | 31.4183 |
| -14 | 27.6253 |
| -13 | 23.8324 |
| -12 | 20.0394 |
| -11 | 16.2464 |
| -10 | 12.4534 |
| -9 | 8.66044 |
| -8 | 4.86746 |
| -7 | 1.07449 |
| -6 | -2.71849 |
| -5 | -6.51147 |

pIDTSMART-AMP:Barcode Btk4

Threshold cycle (Ct)

| Log DNA (g) | Ct |
| --- | --- |
| -9 | 8.90539 |
| -9 | 8.86282 |
| -9 | 8.87053 |
| -10 | 12.5598 |
| -10 | 12.4594 |
| -10 | 12.3736 |
| -11 | 16.2955 |
| -11 | 16.491 |
| -11 | 16.542 |
| -12 | 20.3552 |
| -12 | 20.1222 |
| -12 | 20.2992 |
| -13 | 23.8683 |
| -13 | 23.7403 |
| -13 | 23.6585 |
| -14 | 26.4496 |
| -14 | 26.6313 |
| -14 | 26.6405 |
| -15 | 29.657 |
| -15 | 30.2392 |
| -15 | 29.8417 |
| -16 | 34.6187 |
| -16 | 32.6943 |
| -16 | 33.804 |

| Log DNA (g) | Trend (adj) |
| --- | --- |
| -18 | 40.6573 |
| -17 | 37.1906 |
| -16 | 33.7239 |
| -15 | 30.2572 |
| -14 | 26.7905 |
| -13 | 23.3238 |
| -12 | 19.8572 |
| -11 | 16.3905 |
| -10 | 12.9238 |
| -9 | 9.45707 |
| -8 | 5.99038 |
| -7 | 2.52368 |
| -6 | -0.94301 |
| -5 | -4.4097 |

pIDTSMART-AMP:Barcode Btk6

Threshold cycle (Ct)

| Log DNA (g) | Ct |
| --- | --- |
| -9 | 10.7419 |
| -9 | 10.8492 |
| -9 | 11.018 |
| -10 | 14.167 |
| -10 | 14.2081 |
| -10 | 14.1874 |
| -11 | 17.4703 |
| -11 | 17.5681 |
| -11 | 17.4777 |
| -12 | 21.1948 |
| -12 | 21.3042 |
| -12 | 21.2825 |
| -13 | 25.0155 |
| -13 | 24.9141 |
| -13 | 24.8337 |
| -14 | 28.5175 |
| -14 | 28.5737 |
| -14 | 28.466 |
| -15 | 31.7706 |
| -15 | 31.9819 |
| -15 | 31.8708 |
| -16 | 0 |
| -16 | 0 |
| -16 | 0 |

| Log DNA (g) | Trend (adj) |
| --- | --- |
| -18 | 42.711 |
| -17 | 39.1347 |
| -16 | 35.5585 |
| -15 | 31.9823 |
| -14 | 28.406 |
| -13 | 24.8298 |
| -12 | 21.2536 |
| -11 | 17.6773 |
| -10 | 14.1011 |
| -9 | 10.5249 |
| -8 | 6.94863 |
| -7 | 3.3724 |
| -6 | -0.203839 |
| -5 | -3.78007 |

pIDTSMART-AMP:Barcode Btk5

Threshold cycle (Ct)

| Log DNA (g) | Ct |
| --- | --- |
| -9 | 7.74869 |
| -9 | 7.74288 |
| -9 | 8.61384 |
| -10 | 12.413 |
| -10 | 12.7773 |
| -10 | 12.723 |
| -11 | 17.0189 |
| -11 | 16.8051 |
| -11 | 16.6659 |
| -12 | 20.7961 |
| -12 | 20.794 |
| -12 | 20.899 |
| -13 | 24.4218 |
| -13 | 24.5429 |
| -13 | 24.5289 |
| -14 | 27.6662 |
| -14 | 27.6262 |
| -14 | 27.5191 |
| -15 | 30.9505 |
| -15 | 31.2921 |
| -15 | 30.7955 |
| -16 | 34.3225 |
| -16 | 35.0502 |
| -16 | 34.5132 |

| Log DNA (g) | Trend (adj) |
| --- | --- |
| -18 | 41.9367 |
| -17 | 38.3478 |
| -16 | 34.7588 |
| -15 | 31.1698 |
| -14 | 27.5808 |
| -13 | 23.9919 |
| -12 | 20.4029 |
| -11 | 16.8139 |
| -10 | 13.2249 |
| -9 | 9.63597 |
| -8 | 6.04699 |
| -7 | 2.45802 |
| -6 | -1.13096 |
| -5 | -4.71993 |

pIDTSMART-AMP:Barcode Btk7

Threshold cycle (Ct)

| Log DNA (g) | Ct |
| --- | --- |
| -9 | 7.40559 |
| -9 | 7.51342 |
| -9 | 7.52778 |
| -10 | 11.9094 |
| -10 | 12.2294 |
| -10 | 12.0771 |
| -11 | 15.9389 |
| -11 | 15.9652 |
| -11 | 16.2255 |
| -12 | 20.1133 |
| -12 | 20.1401 |
| -12 | 19.892 |
| -13 | 23.9043 |
| -13 | 23.7389 |
| -13 | 23.681 |
| -14 | 27.2102 |
| -14 | 27.2885 |
| -14 | 27.0669 |
| -15 | 30.8159 |
| -15 | 30.5837 |
| -15 | 31.1245 |
| -16 | 35.0624 |
| -16 | 35.0359 |
| -16 | 35.7444 |

| Log DNA (g) | Trend (adj) |
| --- | --- |
| -18 | 42.4536 |
| -17 | 38.6767 |
| -16 | 34.8998 |
| -15 | 31.1228 |
| -14 | 27.3459 |
| -13 | 23.569 |
| -12 | 19.792 |
| -11 | 16.0151 |
| -10 | 12.2382 |
| -9 | 8.46125 |
| -8 | 4.68432 |
| -7 | 0.907391 |
| -6 | -2.86954 |
| -5 | -6.64647 |

pIDTSMART-AMP:Barcode Btk8

Threshold cycle (Ct)

| Log DNA (g) | Ct |
| --- | --- |
| -9 | 7.97006 |
| -9 | 7.80666 |
| -9 | 7.94316 |
| -10 | 12.0639 |
| -10 | 12.2632 |
| -10 | 12.0792 |
| -11 | 16.3189 |
| -11 | 16.4252 |
| -11 | 16.329 |
| -12 | 19.9709 |
| -12 | 20.2202 |
| -12 | 20.1741 |
| -13 | 23.7419 |
| -13 | 23.82 |
| -13 | 23.6329 |
| -14 | 27.0437 |
| -14 | 27.0201 |
| -14 | 27.0011 |
| -15 | 30.5656 |
| -15 | 30.6855 |
| -15 | 30.958 |
| -16 | 33.7588 |
| -16 | 34.3747 |
| -16 | 33.7421 |

| Log DNA (g) | Trend (adj) |
| --- | --- |
| -18 | 41.3623 |
| -17 | 37.7695 |
| -16 | 34.1767 |
| -15 | 30.5839 |
| -14 | 26.9911 |
| -13 | 23.3984 |
| -12 | 19.8056 |
| -11 | 16.2128 |
| -10 | 12.62 |
| -9 | 9.02724 |
| -8 | 5.43446 |
| -7 | 1.84168 |
| -6 | -1.7511 |
| -5 | -5.34388 |

pIDTSMART-AMP:Barcode Btk9

Threshold cycle (Ct)

| Log DNA (g) | Ct |
| --- | --- |
| -9 | 6.93655 |
| -9 | 6.99809 |
| -9 | 6.92518 |
| -10 | 12.0651 |
| -10 | 11.7894 |
| -10 | 12.0801 |
| -11 | 16.2139 |
| -11 | 15.8617 |
| -11 | 16.1381 |
| -12 | 20.3004 |
| -12 | 20.2959 |
| -12 | 20.6033 |
| -13 | 24.2965 |
| -13 | 24.4123 |
| -13 | 24.2672 |
| -14 | 27.7443 |
| -14 | 27.7796 |
| -14 | 28.0532 |
| -15 | 31.7668 |
| -15 | 31.6118 |
| -15 | 31.4378 |
| -16 | 35.4315 |
| -16 | 36.8337 |
| -16 | 35.4079 |

| Log DNA (g) | Trend (adj) |
| --- | --- |
| -18 | 43.3257 |
| -17 | 39.4498 |
| -16 | 35.5739 |
| -15 | 31.698 |
| -14 | 27.8221 |
| -13 | 23.9462 |
| -12 | 20.0702 |
| -11 | 16.1943 |
| -10 | 12.3184 |
| -9 | 8.44249 |
| -8 | 4.56658 |
| -7 | 0.690664 |
| -6 | -3.18525 |
| -5 | -7.06116 |

pIDTSMART-AMP:Barcode Btk3

Threshold cycle (Ct)

| Log DNA (g) | Ct |
| --- | --- |
| -9 | 10.0988 |
| -9 | 10.2653 |
| -9 | 10.3154 |
| -10 | 13.2127 |
| -10 | 13.3506 |
| -10 | 13.4913 |
| -11 | 16.5144 |
| -11 | 16.4439 |
| -11 | 16.5925 |
| -12 | 20.4637 |
| -12 | 20.4774 |
| -12 | 20.5311 |
| -13 | 23.6756 |
| -13 | 23.6044 |
| -13 | 23.4799 |
| -14 | 26.7908 |
| -14 | 26.6648 |
| -14 | 26.6868 |
| -15 | 30.0449 |
| -15 | 29.9611 |
| -15 | 29.9786 |
| -16 | 33.2267 |
| -16 | 33.6277 |
| -16 | 33.5531 |

| Log DNA (g) | Trend (adj) |
| --- | --- |
| -18 | 40.1924 |
| -17 | 36.8444 |
| -16 | 33.4963 |
| -15 | 30.1483 |
| -14 | 26.8002 |
| -13 | 23.4522 |
| -12 | 20.1041 |
| -11 | 16.7561 |
| -10 | 13.408 |
| -9 | 10.06 |
| -8 | 6.71192 |
| -7 | 3.36387 |
| -6 | 0.0158158 |
| -5 | -3.33224 |

⁂

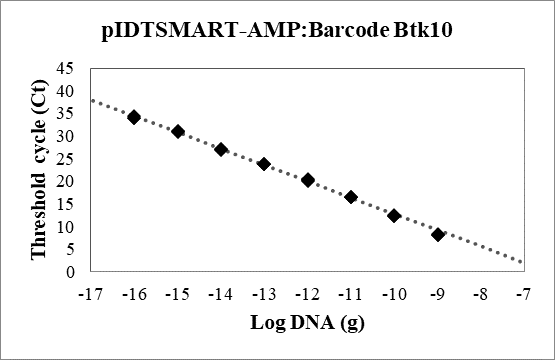

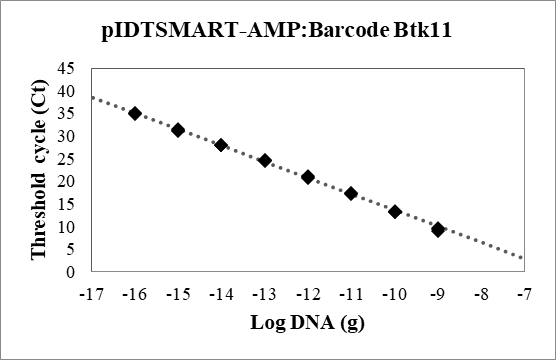

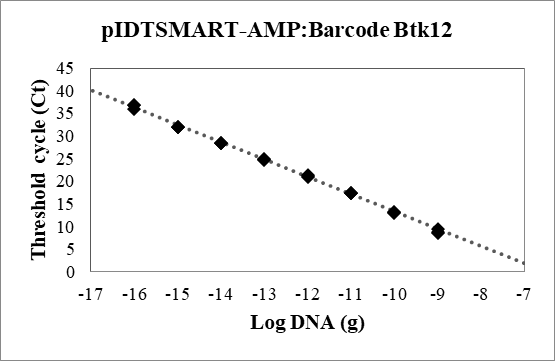

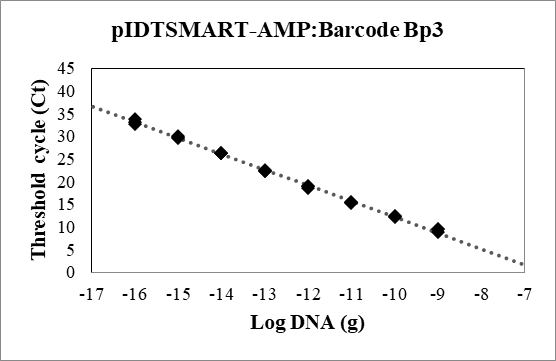

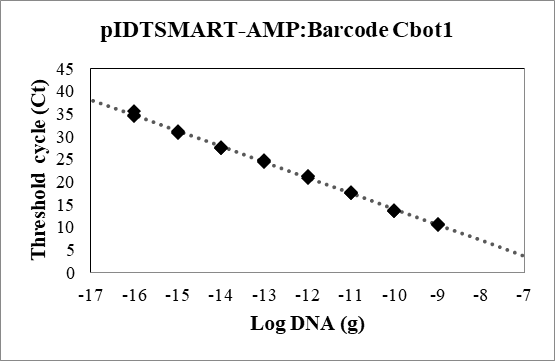

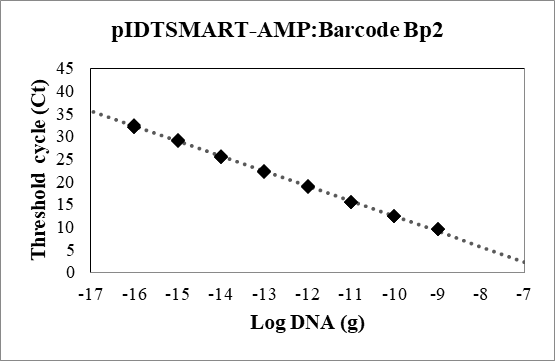

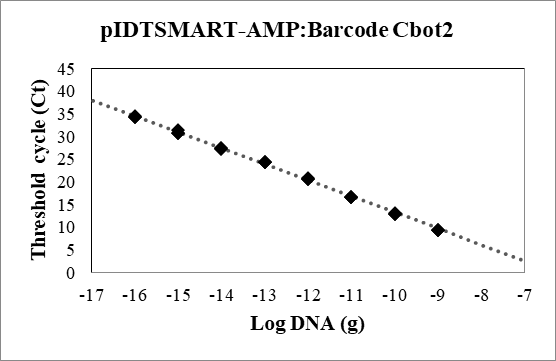

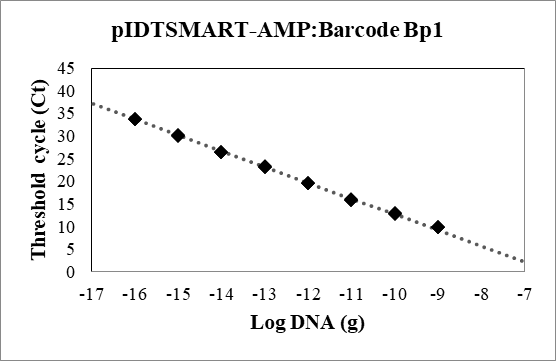

pIDTSMART-AMP:Barcode Btk10

Threshold cycle (Ct)

| Log DNA (g) | Ct |
| --- | --- |
| -9 | 8.16334 |
| -9 | 8.22482 |
| -9 | 8.26724 |
| -10 | 12.3269 |
| -10 | 12.417 |
| -10 | 12.5316 |
| -11 | 16.4578 |
| -11 | 16.5042 |
| -11 | 16.5908 |
| -12 | 20.1116 |
| -12 | 20.3888 |
| -12 | 20.3029 |
| -13 | 23.7924 |
| -13 | 23.7927 |
| -13 | 23.8797 |
| -14 | 27.0357 |
| -14 | 27.0229 |
| -14 | 26.8806 |
| -15 | 30.9817 |
| -15 | 31.0411 |
| -15 | 31.1697 |
| -16 | 34.3074 |
| -16 | 34.0062 |
| -16 | 33.9594 |

| Log DNA (g) | Trend (adj) |
| --- | --- |
| -18 | 41.5412 |
| -17 | 37.9497 |
| -16 | 34.3583 |
| -15 | 30.7668 |
| -14 | 27.1753 |
| -13 | 23.5838 |
| -12 | 19.9924 |
| -11 | 16.4009 |
| -10 | 12.8094 |
| -9 | 9.21793 |
| -8 | 5.62646 |
| -7 | 2.03498 |
| -6 | -1.55649 |
| -5 | -5.14797 |

pIDTSMART-AMP:Barcode Btk11

Threshold cycle (Ct)

| Log DNA (g) | Ct |
| --- | --- |
| -9 | 9.51909 |
| -9 | 9.57253 |
| -9 | 9.00188 |
| -10 | 13.2935 |
| -10 | 13.3541 |
| -10 | 13.223 |
| -11 | 17.2704 |
| -11 | 17.3573 |
| -11 | 17.4015 |
| -12 | 20.9397 |
| -12 | 21.0682 |
| -12 | 20.7538 |
| -13 | 24.7832 |
| -13 | 24.6756 |
| -13 | 24.5303 |
| -14 | 27.9882 |
| -14 | 27.979 |
| -14 | 28.0283 |
| -15 | 31.0546 |
| -15 | 31.4661 |
| -15 | 31.2046 |
| -16 | 35.0422 |
| -16 | 34.8894 |
| -16 | 35.0915 |

| Log DNA (g) | Trend (adj) |
| --- | --- |
| -18 | 42.2125 |
| -17 | 38.6415 |
| -16 | 35.0704 |
| -15 | 31.4994 |
| -14 | 27.9283 |
| -13 | 24.3573 |
| -12 | 20.7863 |
| -11 | 17.2152 |
| -10 | 13.6442 |
| -9 | 10.0731 |
| -8 | 6.50209 |
| -7 | 2.93105 |
| -6 | -0.639996 |
| -5 | -4.21104 |

pIDTSMART-AMP:Barcode Btk12

Threshold cycle (Ct)

| Log DNA (g) | Ct |
| --- | --- |
| -9 | 8.54874 |
| -9 | 8.81177 |
| -9 | 9.38485 |
| -10 | 12.9963 |
| -10 | 13.193 |
| -10 | 13.2904 |
| -11 | 17.351 |
| -11 | 17.4285 |
| -11 | 17.4376 |
| -12 | 21.1577 |
| -12 | 21.2786 |
| -12 | 20.8654 |
| -13 | 24.9761 |
| -13 | 24.972 |
| -13 | 24.6432 |
| -14 | 28.4994 |
| -14 | 28.5272 |
| -14 | 28.4198 |
| -15 | 31.9246 |
| -15 | 31.9739 |
| -15 | 32.0236 |
| -16 | 36.764 |
| -16 | 36.9056 |
| -16 | 35.8639 |

| Log DNA (g) | Trend (adj) |
| --- | --- |
| -18 | 43.9887 |
| -17 | 40.162 |
| -16 | 36.3354 |
| -15 | 32.5088 |
| -14 | 28.6821 |
| -13 | 24.8555 |
| -12 | 21.0289 |
| -11 | 17.2023 |
| -10 | 13.3756 |
| -9 | 9.549 |
| -8 | 5.72237 |
| -7 | 1.89574 |
| -6 | -1.93089 |
| -5 | -5.75752 |

pIDTSMART-AMP:Barcode Bp3

Threshold cycle (Ct)

| Log DNA (g) | Ct |
| --- | --- |
| -9 | 8.94585 |
| -9 | 9.47884 |
| -9 | 9.52441 |
| -10 | 12.4496 |
| -10 | 12.4155 |
| -10 | 12.2906 |
| -11 | 15.4184 |
| -11 | 15.5092 |
| -11 | 15.3506 |
| -12 | 19.0446 |
| -12 | 18.4882 |
| -12 | 19.0178 |
| -13 | 22.49 |
| -13 | 22.3999 |
| -13 | 22.3771 |
| -14 | 26.3498 |
| -14 | 26.3142 |
| -14 | 26.305 |
| -15 | 29.9143 |
| -15 | 29.7153 |
| -15 | 29.9903 |
| -16 | 33.1553 |
| -16 | 33.8921 |
| -16 | 32.7283 |

| Log DNA (g) | Trend (adj) |
| --- | --- |
| -18 | 40.2528 |
| -17 | 36.7347 |
| -16 | 33.2166 |
| -15 | 29.6985 |
| -14 | 26.1803 |
| -13 | 22.6622 |
| -12 | 19.1441 |
| -11 | 15.626 |
| -10 | 12.1078 |
| -9 | 8.58971 |
| -8 | 5.07159 |
| -7 | 1.55346 |
| -6 | -1.96466 |
| -5 | -5.48279 |

pIDTSMART-AMP:Barcode Cbot1

Threshold cycle (Ct)

| Log DNA (g) | Ct |
| --- | --- |
| -9 | 10.4264 |
| -9 | 10.7266 |
| -9 | 10.7341 |
| -10 | 13.5366 |
| -10 | 13.6707 |
| -10 | 13.7358 |
| -11 | 17.8165 |
| -11 | 17.587 |
| -11 | 17.5623 |
| -12 | 21.3338 |
| -12 | 21.3064 |
| -12 | 20.9679 |
| -13 | 24.873 |
| -13 | 24.4893 |
| -13 | 24.7109 |
| -14 | 27.6868 |
| -14 | 27.5712 |
| -14 | 27.572 |
| -15 | 30.9827 |
| -15 | 31.16 |
| -15 | 30.8029 |
| -16 | 34.8112 |
| -16 | 35.5564 |
| -16 | 34.5317 |

| Log DNA (g) | Trend (adj) |
| --- | --- |
| -18 | 41.6056 |
| -17 | 38.1599 |
| -16 | 34.7143 |
| -15 | 31.2686 |
| -14 | 27.8229 |
| -13 | 24.3773 |
| -12 | 20.9316 |
| -11 | 17.4859 |
| -10 | 14.0403 |
| -9 | 10.5946 |
| -8 | 7.14896 |
| -7 | 3.70329 |
| -6 | 0.257628 |
| -5 | -3.18804 |

pIDTSMART-AMP:Barcode Bp2

Threshold cycle (Ct)

| Log DNA (g) | Ct |
| --- | --- |
| -9 | 9.619 |
| -9 | 9.57406 |
| -9 | 9.6245 |
| -10 | 12.4654 |
| -10 | 12.41 |
| -10 | 12.4562 |
| -11 | 15.3988 |
| -11 | 15.4447 |
| -11 | 15.5482 |
| -12 | 18.8774 |
| -12 | 18.9755 |
| -12 | 18.9279 |
| -13 | 22.1373 |
| -13 | 22.3186 |
| -13 | 22.421 |
| -14 | 25.5123 |
| -14 | 25.5584 |
| -14 | 25.5236 |
| -15 | 29.1562 |
| -15 | 28.8801 |
| -15 | 29.2161 |
| -16 | 32.0951 |
| -16 | 32.5957 |
| -16 | 32.1302 |

| Log DNA (g) | Trend (adj) |
| --- | --- |
| -18 | 38.894 |
| -17 | 35.5711 |
| -16 | 32.2483 |
| -15 | 28.9254 |
| -14 | 25.6025 |
| -13 | 22.2796 |
| -12 | 18.9567 |
| -11 | 15.6339 |
| -10 | 12.311 |
| -9 | 8.98808 |
| -8 | 5.6652 |
| -7 | 2.34232 |
| -6 | -0.980565 |
| -5 | -4.30345 |

pIDTSMART-AMP:Barcode Cbot2

Threshold cycle (Ct)

| Log DNA (g) | Ct |
| --- | --- |
| -9 | 9.39805 |
| -9 | 9.39103 |
| -9 | 9.40693 |
| -10 | 12.9413 |
| -10 | 12.8673 |
| -10 | 13.1624 |
| -11 | 16.6285 |
| -11 | 16.6344 |
| -11 | 16.7316 |
| -12 | 20.6754 |
| -12 | 20.9079 |
| -12 | 20.6951 |
| -13 | 24.322 |
| -13 | 24.44 |
| -13 | 24.5055 |
| -14 | 27.4724 |
| -14 | 27.3034 |
| -14 | 27.4627 |
| -15 | 30.7183 |
| -15 | 30.6979 |
| -15 | 31.3677 |
| -16 | 34.2543 |
| -16 | 34.3576 |
| -16 | 34.4921 |

| Log DNA (g) | Trend (adj) |
| --- | --- |
| -18 | 41.6043 |
| -17 | 38.062 |
| -16 | 34.5197 |
| -15 | 30.9774 |
| -14 | 27.4351 |
| -13 | 23.8928 |
| -12 | 20.3505 |
| -11 | 16.8082 |
| -10 | 13.2659 |
| -9 | 9.72355 |
| -8 | 6.18124 |
| -7 | 2.63894 |
| -6 | -0.903365 |
| -5 | -4.44567 |

pIDTSMART-AMP:Barcode Bp1

Threshold cycle (Ct)

| Log DNA (g) | Ct |
| --- | --- |
| -9 | 9.96022 |
| -9 | 9.98536 |
| -9 | 9.84526 |
| -10 | 12.8252 |
| -10 | 13.0251 |
| -10 | 12.9098 |
| -11 | 15.8468 |
| -11 | 15.9401 |
| -11 | 16.0543 |
| -12 | 19.6501 |
| -12 | 19.6718 |
| -12 | 19.5621 |
| -13 | 23.1874 |
| -13 | 23.2651 |
| -13 | 23.1789 |
| -14 | 26.5111 |
| -14 | 26.4587 |
| -14 | 26.5436 |
| -15 | 30.2037 |
| -15 | 30.2913 |
| -15 | 30.0194 |
| -16 | 33.6659 |
| -16 | 33.7891 |
| -16 | 33.8031 |

| Log DNA (g) | Trend (adj) |
| --- | --- |
| -18 | 40.6748 |
| -17 | 37.1739 |
| -16 | 33.6729 |
| -15 | 30.172 |
| -14 | 26.6711 |
| -13 | 23.1702 |
| -12 | 19.6692 |
| -11 | 16.1683 |
| -10 | 12.6674 |
| -9 | 9.16645 |
| -8 | 5.66552 |
| -7 | 2.1646 |
| -6 | -1.33633 |
| -5 | -4.83726 |

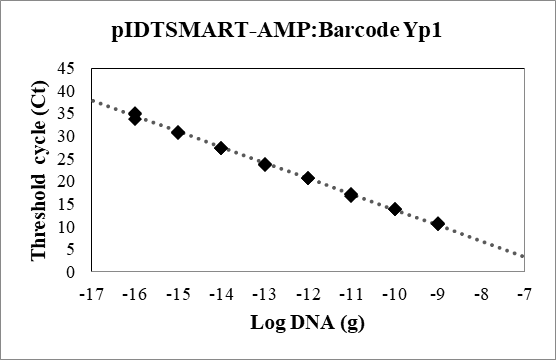

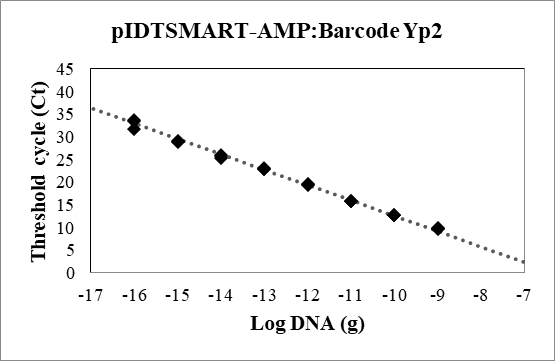

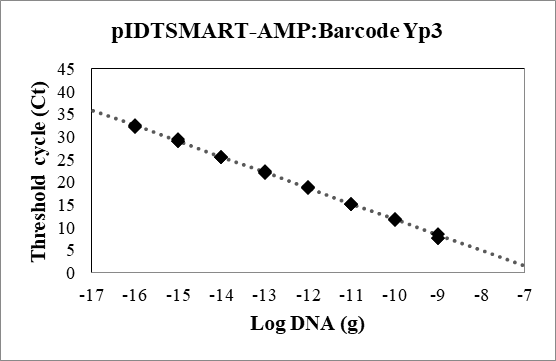

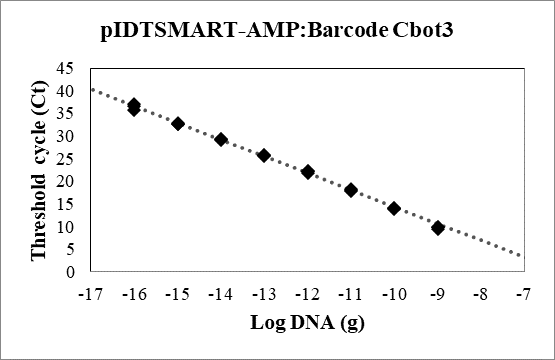

pIDTSMART-AMP:Barcode Yp1

Threshold cycle (Ct)

| Log DNA (g) | Ct |
| --- | --- |
| -9 | 10.6057 |
| -9 | 10.8722 |
| -9 | 10.5544 |
| -10 | 13.7742 |
| -10 | 13.864 |
| -10 | 13.9065 |
| -11 | 17.1283 |
| -11 | 16.8371 |
| -11 | 17.1143 |
| -12 | 20.7912 |
| -12 | 20.7836 |
| -12 | 20.6952 |
| -13 | 23.8011 |
| -13 | 23.8962 |
| -13 | 23.6441 |
| -14 | 27.2767 |
| -14 | 27.4233 |
| -14 | 27.2813 |
| -15 | 30.8155 |
| -15 | 30.7342 |
| -15 | 30.7926 |
| -16 | 34.8677 |
| -16 | 33.7318 |
| -16 | 35.17 |

| Log DNA (g) | Trend (adj) |
| --- | --- |
| -18 | 41.3694 |
| -17 | 37.91 |
| -16 | 34.4505 |
| -15 | 30.991 |
| -14 | 27.5316 |
| -13 | 24.0721 |
| -12 | 20.6126 |
| -11 | 17.1532 |
| -10 | 13.6937 |
| -9 | 10.2342 |
| -8 | 6.77476 |
| -7 | 3.3153 |
| -6 | -0.144169 |
| -5 | -3.60364 |

pIDTSMART-AMP:Barcode Yp2

Threshold cycle (Ct)

| Log DNA (g) | Ct |
| --- | --- |
| -9 | 9.75818 |
| -9 | 9.86518 |
| -9 | 9.89997 |
| -10 | 12.7898 |
| -10 | 12.8024 |
| -10 | 12.8313 |
| -11 | 15.8173 |
| -11 | 15.7441 |
| -11 | 15.789 |
| -12 | 19.4279 |
| -12 | 19.5212 |
| -12 | 19.5752 |
| -13 | 23.1594 |
| -13 | 22.8179 |
| -13 | 22.8856 |
| -14 | 26.0104 |
| -14 | 25.4311 |
| -14 | 25.8178 |
| -15 | 29.1862 |
| -15 | 29.2032 |
| -15 | 28.9755 |
| -16 | 33.8127 |
| -16 | 33.5171 |
| -16 | 31.814 |

| Log DNA (g) | Trend (adj) |
| --- | --- |
| -18 | 39.7947 |
| -17 | 36.3935 |
| -16 | 32.9922 |
| -15 | 29.591 |
| -14 | 26.1897 |
| -13 | 22.7885 |
| -12 | 19.3872 |
| -11 | 15.986 |
| -10 | 12.5847 |
| -9 | 9.18349 |
| -8 | 5.78224 |
| -7 | 2.381 |
| -6 | -1.02025 |
| -5 | -4.4215 |

pIDTSMART-AMP:Barcode Yp3

Threshold cycle (Ct)

| Log DNA (g) | Ct |
| --- | --- |
| -9 | 7.78711 |
| -9 | 8.5692 |
| -9 | 7.76993 |
| -10 | 11.8217 |
| -10 | 11.6077 |
| -10 | 11.9178 |
| -11 | 15.1152 |
| -11 | 15.1343 |
| -11 | 15.2854 |
| -12 | 18.9539 |
| -12 | 18.9522 |
| -12 | 18.7423 |
| -13 | 22.3758 |
| -13 | 22.2498 |
| -13 | 21.9677 |
| -14 | 25.5614 |
| -14 | 25.531 |
| -14 | 25.5005 |
| -15 | 29.1801 |
| -15 | 29.0744 |
| -15 | 29.4944 |
| -16 | 32.536 |
| -16 | 32.436 |
| -16 | 32.1551 |

| Log DNA (g) | Trend (adj) |
| --- | --- |
| -18 | 39.4209 |
| -17 | 35.974 |
| -16 | 32.5271 |
| -15 | 29.0802 |
| -14 | 25.6333 |
| -13 | 22.1864 |
| -12 | 18.7395 |
| -11 | 15.2926 |
| -10 | 11.8457 |
| -9 | 8.39884 |
| -8 | 4.95194 |
| -7 | 1.50504 |
| -6 | -1.94186 |
| -5 | -5.38876 |

pIDTSMART-AMP:Barcode Cbot3

Threshold cycle (Ct)

| Log DNA (g) | Ct |
| --- | --- |
| -9 | 9.86378 |
| -9 | 9.56145 |
| -9 | 9.78801 |
| -10 | 14.0442 |
| -10 | 13.8637 |
| -10 | 14.1239 |
| -11 | 18.1963 |
| -11 | 17.8223 |
| -11 | 17.9753 |
| -12 | 21.7335 |
| -12 | 22.1557 |
| -12 | 22.1525 |
| -13 | 25.6683 |
| -13 | 25.7059 |
| -13 | 25.8748 |
| -14 | 29.3369 |
| -14 | 29.0601 |
| -14 | 29.2514 |
| -15 | 32.5856 |
| -15 | 32.6135 |
| -15 | 32.7678 |
| -16 | 36.5071 |
| -16 | 35.7465 |
| -16 | 37.1018 |

| Log DNA (g) | Trend (adj) |
| --- | --- |
| -18 | 43.9935 |
| -17 | 40.2877 |
| -16 | 36.5818 |
| -15 | 32.876 |
| -14 | 29.1701 |
| -13 | 25.4643 |
| -12 | 21.7584 |
| -11 | 18.0526 |
| -10 | 14.3467 |
| -9 | 10.6409 |
| -8 | 6.93503 |
| -7 | 3.22918 |
| -6 | -0.476669 |
| -5 | -4.18252 |

Figure S1. qPCR standard curves generated for the remainder of the 21 barcodes created by the barCoder algorithm. Template DNA is the indicated barcode in the pIDTSMART-AMP plasmid backbone. For each standard curve, data from three replicates and a trendline are shown.

⁂ Ct value not determinable for 3/3 replicates.
